# High-performance genetically-encoded green and red fluorescent biosensors for pyruvate

**DOI:** 10.1101/2025.04.17.649293

**Authors:** Shosei Imai, Saaya Hario, Carl Suerte, Issei Yamaguchi, Kenryo Sakoi, Adam Thuen, Brekken Meznarich, Mikhail Drobizhev, Takuya Terai, Kei Takahashi-Yamashiro, Robert E. Campbell

## Abstract

Pyruvate is the end-product of glycolysis and a central metabolite involved in many biochemical pathways. However, a lack of high-performance (i.e., Δ*F*/*F*_0_ > 10) single fluorescent protein (FP)-based biosensors has hindered efforts to investigate the physiological role of pyruvate. Here, we present the GreenPy1 and ApplePy1 series, which are green FP (GFP)-based and red FP (RFP)-based pyruvate biosensors, respectively. Both series exhibit large fluorescence intensity change (Δ*F*/*F*_0_ ∼ 20 to >40) and a range of affinities (10s of μM to several mM). We demonstrate the utility of these pyruvate biosensors for multicolor imaging of metabolite concentration changes in mammalian cells.

## Introduction

Pyruvate is an end-product of glycolysis and plays important roles in metabolism. It is converted into ethanol or lactate by fermentation under anaerobic conditions, and fuels the TCA cycle and oxidative phosphorylation in mitochondria under aerobic conditions. Pyruvate is also a substrate for the biosynthesis of many biomolecules including amino acids and fatty acids and regulates several pathological conditions such as cancer and neuronal diseases^1^. Given its many roles, revealing the spatiotemporal dynamics of pyruvate would provide an improved understanding of cellular metabolism.

Fluorescent protein (FP)-based biosensors are some of the most useful and widely used imaging tools, and they have been extensively engineered to provide a wide range of colors and specificities for various target molecules and monatomic ions. These FP-based biosensors can have a variety of different designs^2^. However, of the established designs,single FP-based biosensors composed of circularly permuted (cp)FPs inserted into sensing domains have proved to be among the most versatile and robust^3^. Traditionally, the majority of such single FP-based biosensors have been developed for applications in neuroscience^4,5^, but in recent years there has been increased demand and effort to develop biosensors applicable to other areas of biology, such as cell metabolism. Accordingly, a growing number of biosensors for metabolites have now been reported^6^ including one’s for glucose^7^, citrate^8^, and L-lactate^9,10^.

A number of FP-based pyruvate biosensors have previously been reported, including several Förster resonance energy transfer (FRET)-based biosensors^11–13^, and the single GFP-based biosensors PyronicSF^14^ and Green Pegassos^15^. The reported FRET-based biosensors have Δ*R*/*R*_0_ = 0.2 ∼ 0.6, PyronicSF has a Δ*F*/*F*_0_ ∼ 2.5 and an apparent *K*_d_ of 480 μM, and Green Pegassos has a Δ*F*/*F*_0_ ∼ 2.3 and an apparent *K*_d_ of 70 μM, as measured *in vitro*. These biosensors have successfully been used to visualize pyruvate dynamics under a variety of conditions. However, the FRET-based ones are generally not practical for multiplexed imaging applications and are limited by their small ratiometric responses. Neither of the two previously reported single FP-based biosensors can be described as being high-performance (i.e., Δ*F*/*F*_0_ > 10) by the current standards of the field, and together they provide limited options for visualizing pyruvate across a wide concentration range.

To overcome the limitations of the previously reported pyruvate biosensors, here we report high-performance green fluorescent (the GreenPy series) and red fluorescent (the ApplePy series) pyruvate biosensors with affinities that span the physiological concentration range (**Fig. 1A**). These biosensors were developed through an extensive process of protein engineering using directed evolution and structure-guided mutagenesis. We demonstrate the utility of the biosensors for imaging of pyruvate concentration changes in mammalian cells.

**Figure 1.**
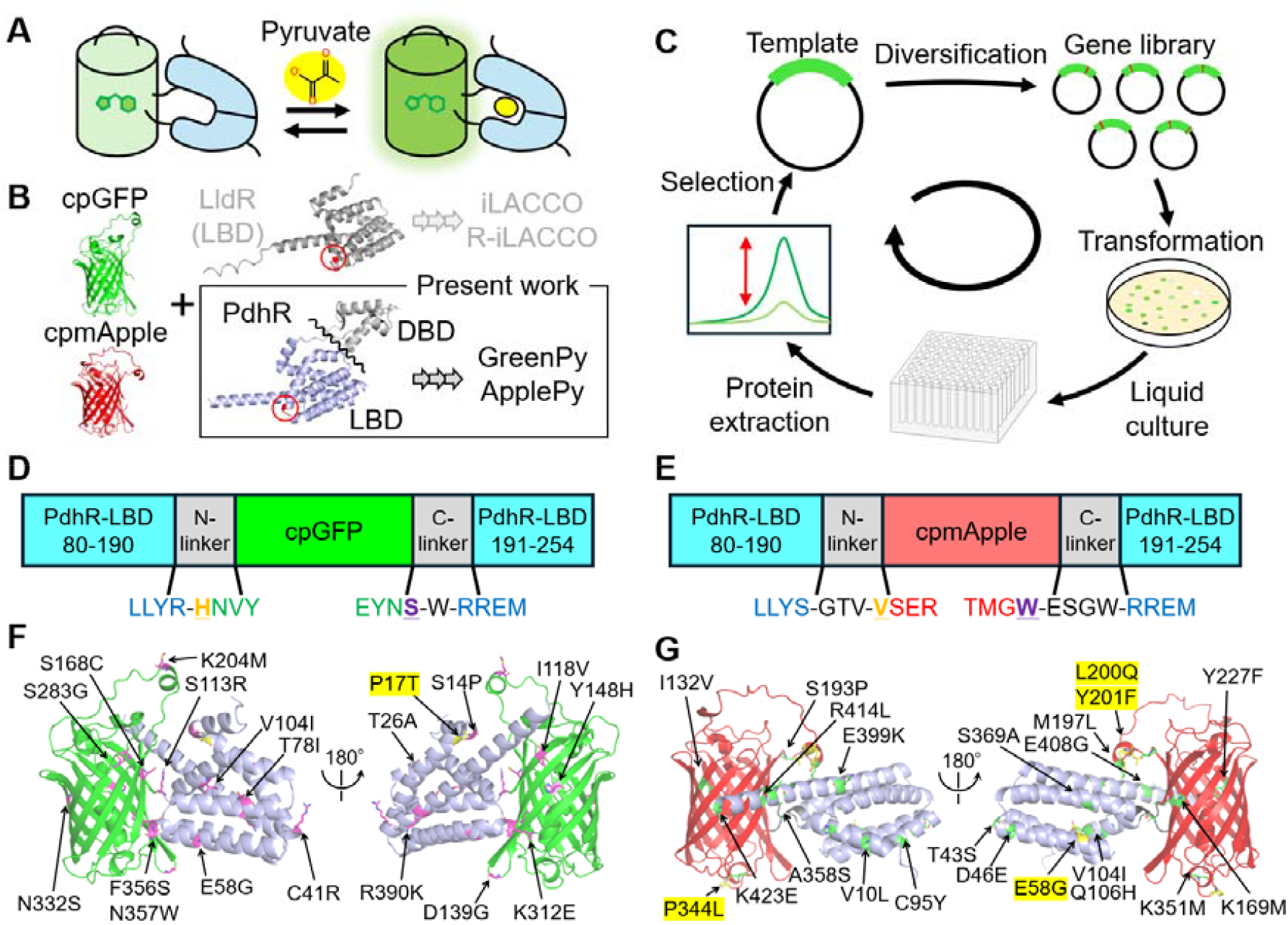
Development of GreenPy and ApplePy. **A**, Schematic representation of PdhR-based GreenPy and ApplePy. **B**, Overall strategy to identify the initial prototype. Red circles indicate the position to insert the cpFPs. **C**, Schematic representation of directed evolution. **D,E**, Schematic representation of the domain structure of GreenPy variants (**D**) and ApplePy variants (**E**). The two gate-post residues^3^ in cpFPs are highlighted in orange and purple and underlined. **F,G**, Structures of GreenPy1High (**F**) and ApplePy1High (**G**) as predicted by AlphaFold3^18^. The mutations relative to GreenPy0.2 and ApplePy0.3 are shown in magenta and green, respectively, and the mutations of other affinity variants are colored yellow.

## Results

### Development of a green pyruvate biosensor

To develop pyruvate biosensors, we took inspiration from our previous work to develop L-lactate biosensors by inserting cpFPs into LldR, an L-lactate-binding transcription factor^9,10^. PdhR is a pyruvate-binding transcription factor that belongs to the same protein subfamily as LldR (FadR subfamily)^16,17^ and therefore seemed like a suitable binding domain for creating pyruvate-specific biosensors. Like all members of the FadR family, PdhR is composed of a ligand binding domain (LBD) and a DNA binding domain (DBD). To construct an initial prototype, we deleted the DBD and inserted cpFPs into the LBD of PdhR. To determine the optimal insertion site, we turned our attention to the similarity between PdhR and LldR, which was previously used to develop the iLACCO series^9^. Their predicted structures^18^ were superimposed and the residues of PdhR (S190 and R191; numbering based on full length PdhR) that correspond to the LldR insertion site used for iLACCO (L187 and V188) were identified (**Fig. 1B**). PyronicSF is based on insertion of cpGFP between residues 188 and 189 of PdhR^14^. When cpGFP derived from iGluSnFR^19^ was inserted into the LBD of PdhR between S190 and R191, the resulting protein exhibited an inverted response of Δ*F*/*F*_0_ = -0.20 in response to 10 mM pyruvate. This prototype was designated as GreenPy0.1 and served as a template for further improvement.

To improve the brightness and increase the Δ*F*/*F*_0_ of the GreenPy0.1 prototype, we used a combination of site-directed mutagenesis and directed evolution. Directed evolution was conducted by screening libraries, generated by site-saturation mutagenesis or random mutagenesis, for variants with improved brightness and pyruvate-dependent fluorescence intensity changes (**Fig. 1C**). We first optimized the linkers that connect the cpFP to the PhdR domain. The initial prototype had a PdhR to cpFP linker (N-linker) with a sequence of DWS and a cpFP to PdhR linker (C-linker) with a sequence of NNV. We systematically tested variants with partial or complete linker deletions (**Supplementary Fig. 1A**). We determined that the variant with a deleted N-linker and a shortened C-linker (NNV→N) exhibited the largest direct response fluorescence change. This variant was designated as GreenPy0.2. From this point on, residues are numbered according to the GreenPy sequence, as provided in **Supplementary Fig. 2**.

Next, we optimized the residues located at or near the linker regions by screening variants in which pairs of residues were fully randomized. For the N-linker, we randomized S113 (equivalent to S190 of PhdR) and the immediately adjacent residue of cpGFP (H114) which has been previously defined as one of two critical gatepost residues^3^. For the C-linker we randomized cpFP gatepost residue F356 and linker residue N357. These efforts yielded a variant with Δ*F*/*F*_0_ ∼ 4.5 but with only ∼ 10% of the brightness of GreenPy0.1. To improve the brightness, we performed two rounds of directed evolution with random mutagenesis of the whole gene using error-prone PCR, followed by a second round of linker optimization. Together, the four rounds of directed evolution resulted in GreenPy0.3, which harbored five mutations (C41R, V104I, S113R, F356S and N357W) and exhibited a Δ*F*/*F*_0_ of 2.0. To further improve both response and brightness, we performed 8 additional rounds of directed evolution and obtained GreenPy0.9, which exhibited Δ*F*/*F*_0_ = 28.5, an apparent *K*_d_ of ∼ 200 μM, and contained 11 point mutations relative to GreenPy0.3 (**Fig. 1D,F** and **Supplementary Fig. 2**).

### Development of a red pyruvate biosensor

To identify an initial prototype of a red fluorescent biosensor for pyruvate, cpmApple derived from R-iLACCO2 (an L-lactate biosensor developed by Yuki Kamijo and Yusuke Nasu from R-iLACCO1^10^), was inserted into the same site of the PdhR LBD as used for GreenPy. The resulting prototype, ApplePy0.1, exhibited Δ*F*/*F*_0_ = -0.11. The initial linkers derived from R-iLACCO2 consisted of an N-linker of DW and a C-linker of EATR. For the optimization of the N-linker, a library with linker lengths ranging from 0 to 3 residues was created and screened (**Supplementary Fig. 1B**). For the optimization of the C-linker, only the third and fourth residues (TR) were randomized because the first two residues (EA) tend to be conserved in cpmApple-based biosensors^3^. The linker-optimized variant, ApplePy0.3, had an N-linker of GTV, a C-linker of EAGW, and a Δ*F*/*F*_0_ of 1.1.

We next conducted directed evolution to further improve the performance. Briefly, this involved randomization of the entire gene, combined with site-directed and site-saturation mutagenesis at the sites of beneficial mutations (see **Supplementary Note 1** for details). This effort eventually led to the discovery of ApplePy0.9 with Δ*F*/*F*_0_ = 17.2, an apparent *K*_d_ of ∼ 700 μM, and 15 point mutations relative to ApplePy0.3 (**Fig. 1E,G** and **Supplementary Fig. 3**).

### Affinity tuning of GreenPy and ApplePy variants

As the concentration of pyruvate *in vivo* has been reported to range from tens to hundreds of µM depending on the organelle and tissue type^14,20,21^, and can reach higher concentrations in certain metabolic disorders^22–24^, we determined that it is important to have variants with affinities that cover the entire physiological concentration range. Towards this goal, we performed several additional rounds of site-directed and random mutagenesis using lower concentrations of pyruvate for the screening assay to identify higher affinity variants, and higher concentrations of pyruvate to identify lower affinity variants. We identified four GreenPy variants that we designated as GreenPy1Lowest, GreenPy1Low, GreenPy1High and GreenPy1Highest (**Supplementary Note 2**). Similarly, we tuned the affinity of ApplePy to develop four corresponding affinity variants (**Supplementary Note 2**). Their apparent *K*_d_ values span from tens of µM to several mM, which covers the entire physiologically relevant pyruvate concentration range.

Among the mutations that affected the affinity for pyruvate, only the sidechain of residue 104 (that is, the position of the V104I mutation which increased the affinity) is predicted to point toward the putative binding cavity (**Fig. 1F,G**). The location of the putative binding cavity is identified by superimposing the model structure with a previously reported crystal structure of NanR, which is also a member of the FadR subfamily^25^ (**Supplementary Fig. 4**). As the sidechains of Ile and Val have similar hydrophobicity but different structures, we suggest that the V104I mutation alters the shape of the binding pocket to better accommodate pyruvate. Other mutations that affected the affinities were introduced into the α-helices of the PdhR. Although they are not predicted to interact directly with a pyruvate, they may play an important role in interactions between the α-helices, indirectly determining the pocket size. Further insight will likely require the experimentally determined structure of the bound state.

In addition to these affinity variants, we also developed control biosensors that are unresponsive to pyruvate but sustain similar fluorescence intensity and exhibit similar pH sensitivity. The control biosensors dGreenPy and dApplePy were engineered by inserting a single glycine residue into the C-linker of GreenPy1High and ApplyPy1High, respectively (**Supplementary Fig. 5**).

### *In vitro* characterization of GreenPy and ApplePy variants

The photophysical properties of the GreenPy variants, as well as the previously reported PyronicSF^14^ and Green Pegassos^15^ biosensors, were determined *in vitro* using purified proteins. The GreenPy1 series exhibited emission intensiometric responses of Δ*F*/*F*_0_ = 20.1, 44.3, 41.0 and 48.2, and affinities of *K*_d,app_ = 35, 200, 370 and 2730 µM, for Highest, High, Low, and Lowest, respectively (**Fig 2A,C**). The dGreenPy control exhibited a Δ*F*/*F*_0_ ∼ 0.1 in going from 0 to 100 mM pyruvate (**Fig 2A**). Since the four affinity variants showed similar values for other photophysical parameters, the results of GreenPy1High are discussed in the main text and the results for the other three variants are presented in **Supplementary Fig. 6** and **Table 1**. The results for PyronicSF and Green Pegassos are shown in **Supplementary Fig. 7** and **Table 1**. Our characterization provided values that were generally in good agreement with the previously published values. For PyronicSF, we found Δ*F*/*F*_0_ = 1.5 (previously 2.5) and an apparent *K*_d_ of ∼ 1910 μM (previously 480 μM). For Green Pegassos we found Δ*F*/*F*_0_ = 0.9 (previously 2.3) and an apparent *K*_d_ of ∼ 62 μM (previously 70 μM).

**Figure 2.**
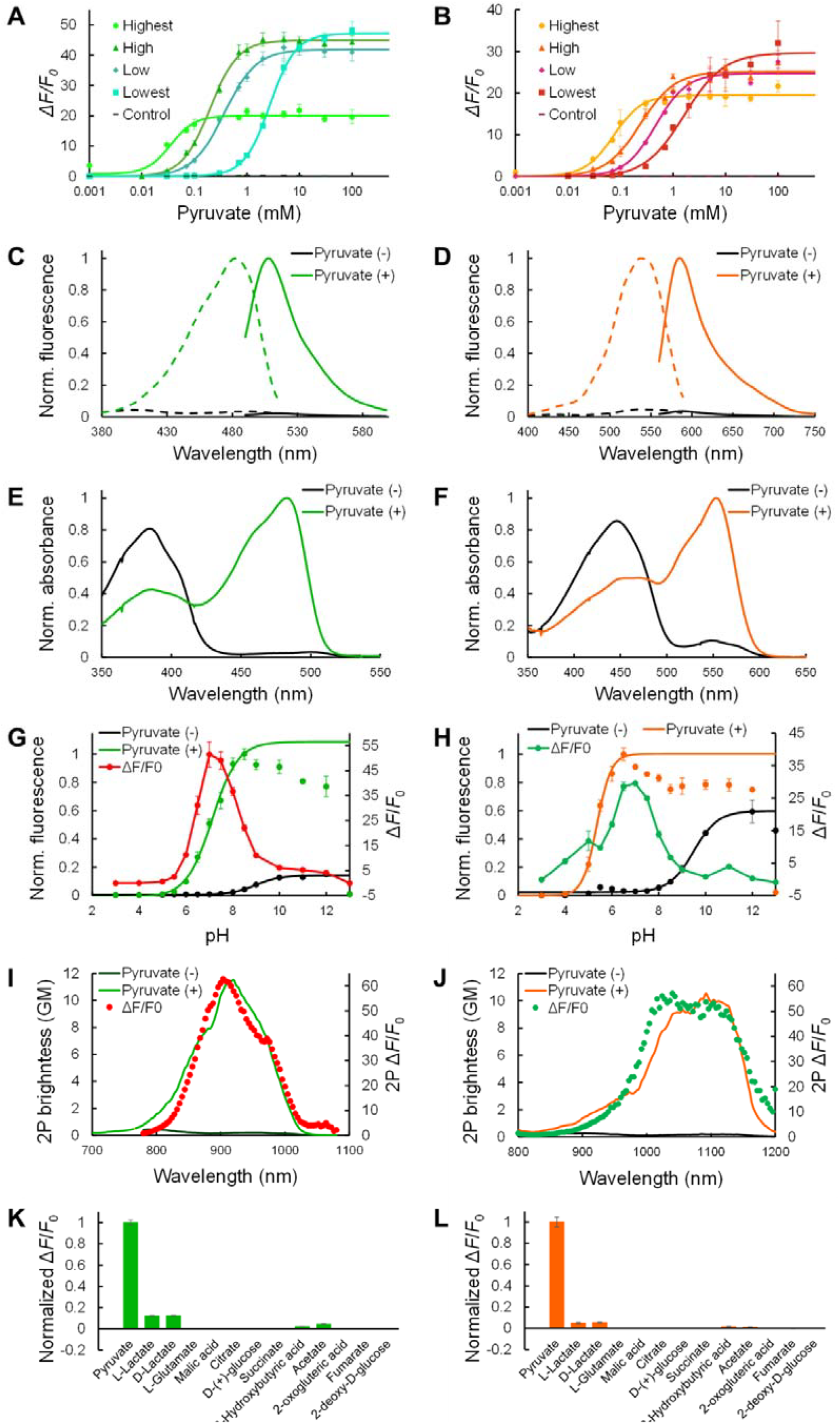
*In vitro* characterization. **A,B**, Pyruvate titration curve of purified GreenPy (**A**) and ApplePy (**B**) variants (n = 3 replicates; mean ± s.d.). **C,D**, Excitation (emission at 555 nm, 640 nm) and emission spectra (excitation at 450 nm, 510 nm) of GreenPy1High (**C**) and ApplePy1High (**D**) in the presence (100 mM) and the absence of pyruvate. **E,F**, Absorbance spectra of GreenPy1High (**E**) and ApplePy1High (**F**) in the presence (10 mM) and absence of pyruvate. **G,H**, pH titration curve of GreenPy1High (**G**) and ApplePy1High (**H**) in the presence (100 mM) and the absence of pyruvate (n = 3 replicates; mean ± s.d.). The curve was fitted to calculate the physiological p*K*_a_ value. **I,J**, Two-photon excitation spectra of GreenPy1High (**I**) and ApplePy1High (**J**) in the presence (10 mM) and absence of pyruvate. 2P, Two-photon. GM, Goeppert-Mayer units. **K,L**, Fluorescence responses of GreenPy1High (**K**) and ApplePy1High (**L**) to various metabolites at 10 mM (n = 3 replicates; mean ± s.d.).

**Table 1.**
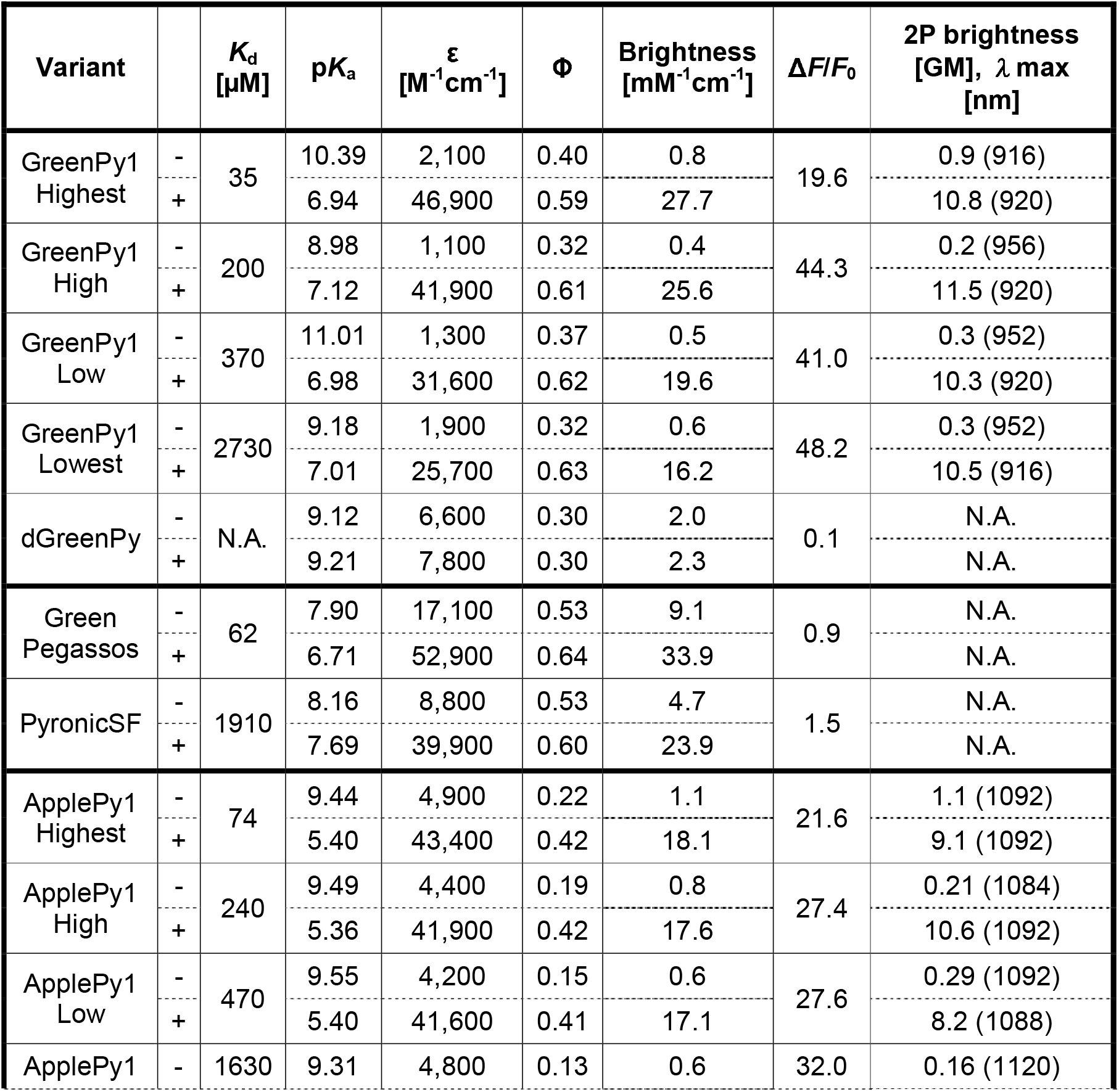
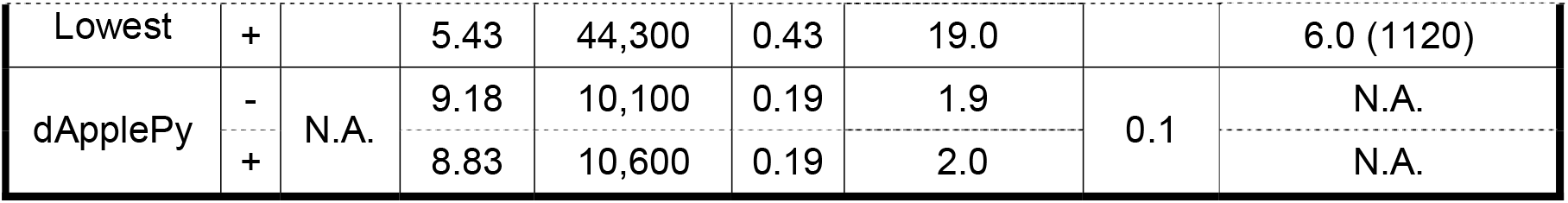
Photophysical and biochemical properties of the green and red pyruvate biosensors.

For GreenPy1High, the absorbance spectrum changes in a ratiometric manner upon binding with pyruvate, with the peak at 384 nm (dim protonated state of the chromophore) decreasing and the peak at 483 nm (bright anionic state of the chromophore) increasing (**Fig 2E**). The effective extinction coefficient at 483 nm increases from 1,100 M^-1^ cm^-1^ in the free state to 41,900 M^-1^ cm^-1^ in the pyruvate-bound state, and the quantum yield increases from 0.32 to 0.61 (**Table 1**). The effective extinction coefficient represents a product of the molar extinction coefficient ε_A_ of the anionic state of chromophore and the fractional concentration of anionic state ρ_A_. In principle, both ε_A_ and ρ_A_ can change upon binding ligand (see **Characterization of two-photon absorption properties** in **Methods** for details). GreenPy1High has an apparent p*K*_a_ of 8.98 in the absence of pyruvate, shifting to 7.12 in the presence of pyruvate (**Fig 2G**), This shift suggests the change of fractional concentration of anionic state at neutral pH upon binding pyruvate, consistent with the observed absorbance change. The analysis of the factors contributing to the fluorescence change of anionic state, namely extinction coefficient, absorption peak position, fractional concentration, and fluorescence quantum yield, for all 4 green biosensors is presented in **Supplementary Table 1**. The p*K*_a_ value of dGreenPy is 9.21 and 9.23 in the pyruvate-unbound and -bound state, respectively. The molecular brightness of GreenPy1High in the pyruvate-bound state is 76% that of EGFP, 107% that of PyronicSF and 75% that of Green Pegassos. Consistent with the absorbance spectrum, the excitation maximum in the presence of pyruvate is observed at 482 nm, with an emission maximum at 508 nm. With two-photon (2P) excitation, GreenPy1High has an excitation peak at 920 nm in the presence of pyruvate with 2P brightness of 11.5 GM (**Fig 2I**). The Δ*F*/*F*_0_ calculated using 2P brightness exceeded 60 between 880 and 930 nm.

The photophysical properties of ApplePy variants were also characterized *in vitro* as purified proteins, and the results of ApplePy1High are discussed here (results for the other three variants are shown in **Supplementary Fig. 8** and **Table 1**). The purified proteins of four affinity variants exhibited emission intensiometric Δ*F*/*F*_0_ = 21.6, 27.4, 27.6 and 32.0, and *K*_d, app_ = 74, 240, 470 and 1630 µM, for Highest, High, Low, and Lowest, respectively (**Fig 2B,D**). The response of dApplePy was negligible (**Fig 2B**). The absorbance change upon binding with pyruvate was found to be ratiometric, with the peak for the protonated state of the chromophore at 446 nm decreasing and the peak for the anionic state of the chromophore at 554 nm increasing (**Fig 2F**). The effective extinction coefficient at 554 nm is 4,400 M^-1^ cm^-1^ in the free state and 41,900 M^-1^ cm^-1^ in the pyruvate-bound state, and the quantum yield increased from 0.19 to 0.42. **Supplementary Table 2** presents the analysis of the factors contributing to the fluorescence change of anionic state, for all 4 red biosensors. The molecular brightness of the pyruvate-bound state is 48% that of mApple and 117% that of R-GECO1. The excitation and emission maxima were 538 nm and 584 nm, respectively. The p*K*_a_ value of ApplePy1High changes from 9.49 to 5.22 upon binding with pyruvate (**Fig 2H**). With 2P excitation, ApplePy1High has an excitation peak at 1092 nm in the presence of pyruvate and a 2P brightness of 10.6 GM (**Fig 2J**). It exhibits a Δ*F*/*F*_0_ of greater than 50 between 1,000 and 1,100 nm.

To determine the specificity for pyruvate, all of the GreenPy1 and ApplePy1 variants were incubated with 10 mM of various metabolites. At 10 mM, we found that the lower affinity variants had greater specificity for pyruvate than the higher affinity variants.

Specifically, the higher affinity variants exhibited substantial responses to lactate, 3-hydroxybutyric acid and acetate (**Fig. 2K** and **Supplementary Fig. 9**). Comparing GreenPy1Lowest (apparent *K*_d_ = 2730 μM) with PyronicSF (apparent *K*_d_ = 1910 μM), and GreenPy1Higest (apparent *K*_d_ = 35 μM) with Green Pegassos (apparent *K*_d_ = 62 μM), which have similar affinities, revealed that GreenPy1 variants are more selective for pyruvate (compare **Supplementary Fig. 6M,O** with **Supplementary Fig. 7I,J**). ApplePy1 variants showed similar results to GreenPy1 variants (**Fig. 2L, Supplementary Fig. 8**).

### Mammalian cell imaging with the GreenPy1 and ApplePy1 series

To determine if the GreenPy1 series and the ApplePy1 series could enable imaging of changes in pyruvate concentration in mammalian cells imaging, we transfected HeLa cells with plasmids encoding each of the eight biosensors and two controls. Addition of pyruvate to a final concentration of 10 mM in the imaging buffer, without any other pharmacological treatment, increased the fluorescence intensity of the GreenPy1 and the ApplePy1 series as shown in **Fig. 3A,B**. For both series, the responses followed the general trend that the lower affinity biosensors gave the largest Δ*F*/*F*_0_, and the higher affinity biosensors gave the smallest Δ*F*/*F*_0_. Assuming that the apparent *K*_d_ values (**Table 1**) are valid for the biosensors in the intracellular milieu, the observed fluorescent changes are consistent with a resting pyruvate concentration in the 10s of μM, rising to several mM with 10 mM extracellular pyruvate.

**Figure 3.**
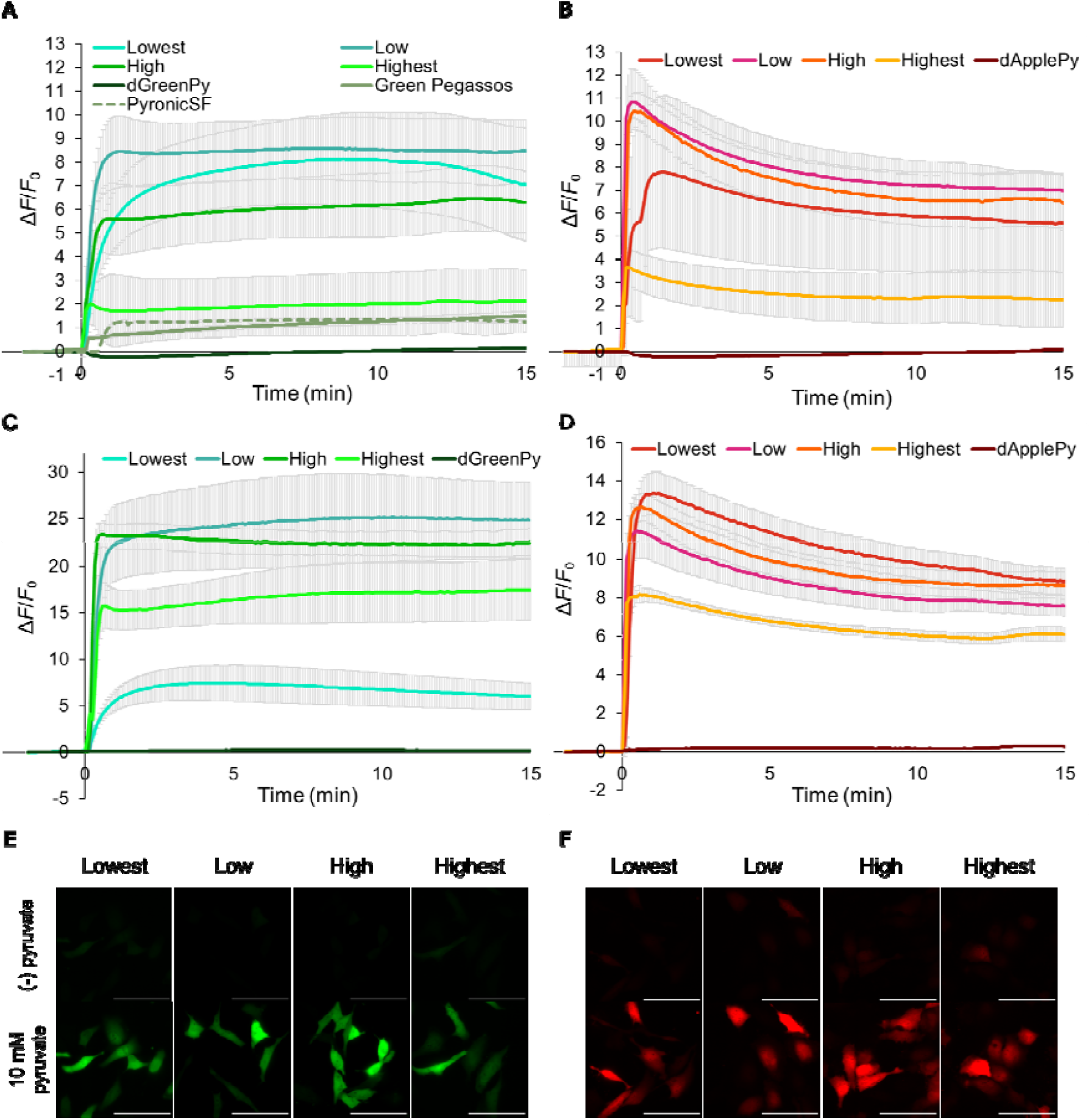
Cell-based characterization of the GreenPy1 series and ApplePy1 series. **A,B**, Time courses of Δ*F*/*F*_0_ of GreenPy1Lowest (n = 12), GreenPy1Low (n = 10), GreenPy1High (n = 10), GreenPy1Highest (n = 9), dGreenPy1 (n = 12), Green Pegassos (n = 8), and PyronicSF (n = 4) (**A**), and ApplePy1Lowest (n = 16), ApplePy1Low (n = 14), ApplePy1High (n = 7), ApplePy1Highest (n = 12), and dApplePy1 (n = 11) (**B**), expressed in HeLa cells that were treated with sodium pyruvate (final concentration of 10 mM). (mean ± s.d.). **C,D**, Time courses of Δ*F*/*F*_0_ of GreenPy1Lowest (n = 15), GreenPy1Low (n = 12), GreenPy1High (n = 11), GreenPy1Highest (n = 7), and dGreenPy1 (n = 9) (**C**), and ApplePy1Lowest (n = 9), ApplePy1Low (n = 6), ApplePy1High (n = 7), ApplePy1Highest (n = 9), and dApplePy1 (n = 4) (**D**), expressed in HeLa cells (pre-treated with iodoacetic acid, nigericin, and rotenone), that were treated as in **A,B**. (mean ± s.d.). **E,F**, Snapshots of before and after the addition of exogenous 10 mM pyruvate to the pre-treated HeLa cells expressing the GreenPy1 series (**E**) or the ApplePy1 series (**F**). Scale bars represent 100 μm.

At the resting pyruvate concentration, the Highest biosensors should exist primarily in the bound state, which is consistent with their relatively small responses upon treatment of the cells with pyruvate. To better evaluate the responsiveness of the Highest biosensors, we decreased the concentration of intracellular pyruvate using a previously described method^9,26^. Briefly, HeLa cells were preincubated with iodoacetic acid to inhibit glycolysis, and the imaging buffer contained nigericin to clamp the pH and rotenone to block mitochondrial metabolism. Under this condition, GreenPy1Highest and ApplePy1Highest variants had much larger Δ*F*/*F*_0_ values compared to the values measured in HeLa cells without pre-treatment, upon treatment with 10 mM extracellular pyruvate (**Fig. 3C-F**).

### Imaging of cytosolic and mitochondrial pyruvate dynamics

To further investigate the use of the GreenPy1 and ApplePy1 series for imaging of pyruvate concentration changes in cytosol and mitochondria, we carried out cell imaging during several pharmacological and other treatments. AR-C155858 is a specific inhibitor of proton-coupled monocarboxylate transporters 1 and 2 (MCT1, MCT2)^27^, which reversibly transport pyruvate and lactic acid across the plasma membrane (**Fig. 4A**). We found that treatment of HeLa cells (expressing the pyruvate biosensors) with AR-C155858 increased the fluorescence intensity of biosensors (**Fig. 4B,C**), consistent with an increase in the cytosolic pyruvate concentration.

**Figure 4.**
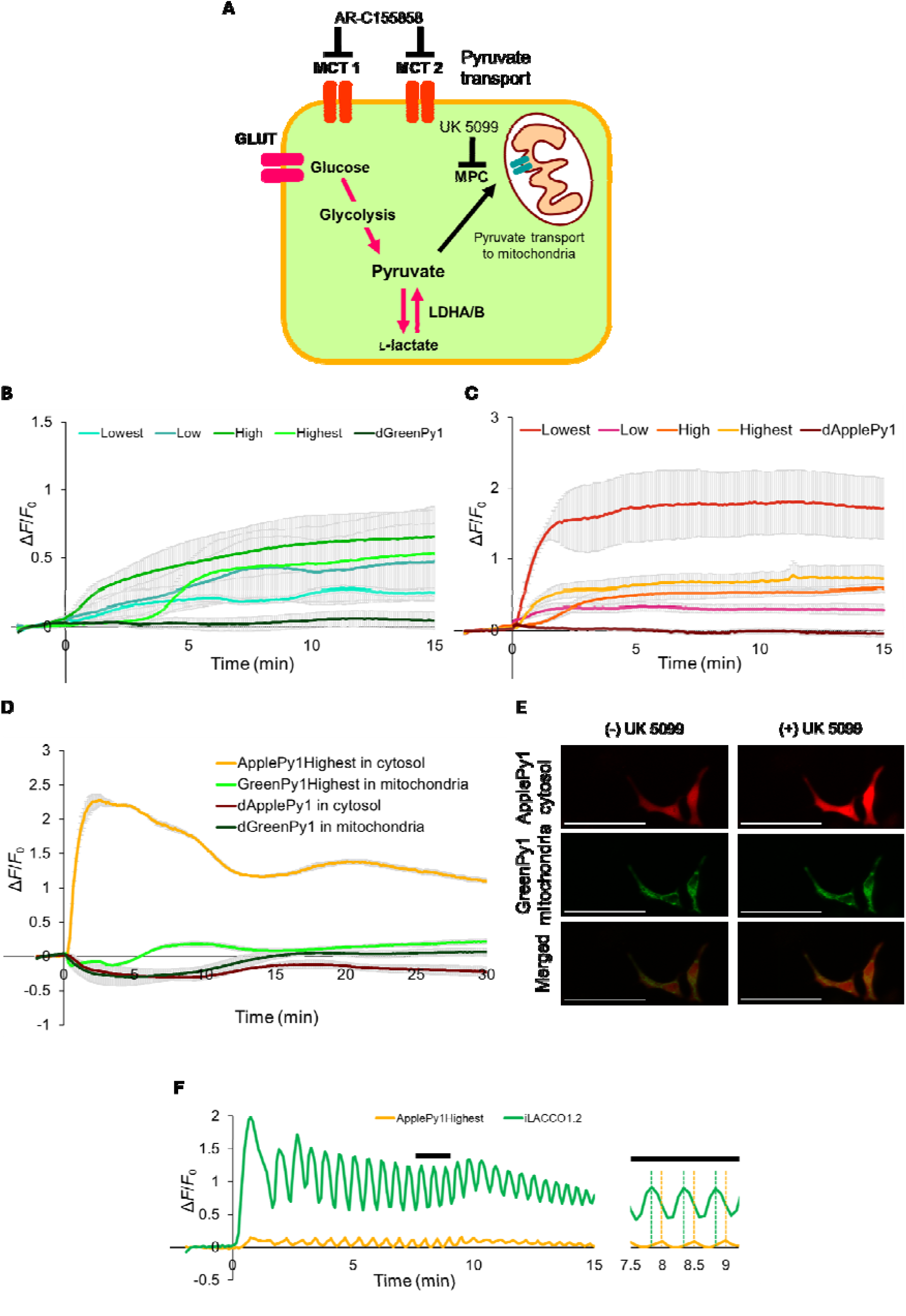
Imaging of cytosolic and mitochondrial pyruvate dynamics. **A**, A schematic of pyruvate dynamics in mammalian cells and inhibitors used in these experiments. **B,C**, Time courses of Δ*F*/*F*_0_ of GreenPy1Lowest (n = 8), GreenPy1Low (n = 5), GreenPy1High (n = 14), GreenPy1Highest (n = 5), and dGreenPy1 (n = 19) (**B**) and ApplePy1Lowest (n = 6), ApplePy1Low (n = 4), ApplePy1High (n = 5), ApplePy1Highest (n = 4), and dApplePy1 (n = 9) (**C**) expressed in cytosol in HeLa cells upon the addition of MCT1,2 inhibitor, AR-C155858 (final concentration of 1 μM). (mean ± s.d.). **D**, Time courses of Δ*F*/*F*_0_ of ApplePy1Highest (n = 2) and dApplePy1 (n = 4) expressed in the cytosol, and GreenPy1Highest (n = 2) and dGreenPy1 (n = 4) expressed in mitochondria, upon the addition of MPC inhibitor, UK 5099 (final concentration of 25 μM). (mean ± s.d.). A set of ApplePy1Highest and GreenPy1Highest and dApplePy1 and dGreenPy1 were expressed in each experiment. **E**, Snapshots of HeLa cells expressing ApplePy1Highest in the cytosol and GreenPy1Highest in mitochondria before and after the addition of UK 5099. **F**, Time courses of Δ*F*/*F*_0_ of ApplePy1Highest and iLACCO1.2 co-expressed in cytosol in starved HeLa cells upon the addition of D-glucose (final concentration of 5 mM). Representative data from one cell is shown in the graph. The right graph showed the enlarged graph from 7.5 - 9 min of left graph (black bar).

Next, to investigate pyruvate transport between the cytosol and mitochondria, we tested the effect of the mitochondrial pyruvate carrier (MPC) inhibitor, UK 5099 (**Fig. 4A**). Cytosolic pyruvate is imported into mitochondria through voltage-dependent anion channel (VDAC) at the outer mitochondrial membrane and MPC at the inner mitochondrial membrane. We co-expressed ApplePy1Highest (with no targeting sequence) and GreenPy1Highest (with 2× COX8) in the cytosol and mitochondria, respectively (**Fig. 4D,E**). The fluorescence of ApplePy1Highest in cytosol increased immediately after the addition of the MPC inhibitor, followed by a gradual decrease. In contrast, GreenPy1Highest did not show a large change in fluorescence. These results imply that inhibition of import of pyruvate into mitochondria did not cause a substantial decrease in the mitochondrial pyruvate concentration.

We have previously visualized oscillations in L-lactate concentration upon the addition of D-glucose to starved mammalian cells^9^. To determine if we could visualize corresponding oscillations in the pyruvate concentration, we co-expressed both iLACCO1.2, a high affinity L-lactate biosensor^9^, and ApplePy1Highest (**Fig. 4F**). ApplePy1Highest responds to high concentrations (> 10 mM) of L-lactate (**Supplementary Fig. 9B**), but this concentration is far greater than the apparent *K*_d_s of both ApplePy1Highest for pyruvate (74 μM) and iLACCO1.2 for lactate (17 μM). We observed that the oscillations of iLACCO1.2 and ApplePy1Highest have a time lag between their phases consistent with each biosensor responding selectively to its respective target.

## Discussion

We have developed a set of genetically encoded green and red biosensors for pyruvate, designated as GreenPy1 and ApplePy1 series. Through extensive directed evolution and affinity tuning, we obtained pyruvate biosensors with high response and brightness. Compared with previously reported FRET-based and single-GFP based pyruvate biosensors, the GreenPy1 series exhibited substantially higher responses and a range of affinities. In addition, the ApplePy1 series is not only comparable to the GreenPy series in terms of performance and versatility, but is also the first reported example of red fluorescent pyruvate biosensor. The red fluorescent ApplePy1 series creates new opportunities for multi-color imaging with existing green fluorescent biosensors, and may enable deeper *in vivo* imaging due to the increased transparency of tissues to longer wavelength light. Both the GreenPy1 and ApplePy1 series showed stable expression and robust responses to changes in pyruvate concentration in mammalian cells.

Building on our previous work to use the LldR L-lactate binding protein to develop high performance L-lactate biosensors^9^, in this work we used PdhR. LldR and PdhR are protein homologs that belong to the FadR subfamily of transcription factors. While periplasmic binding proteins (PBPs)^19,28,29^ and G-protein-coupled receptors (GPCRs)^30–32^ have been frequently used to develop biosensors for a wide range of targets, the use of transcription factors has been relatively unexplored. Together with our previous efforts to develop LldR-based L-lactate biosensors^9,10^, this work convincingly demonstrates the utility of FadR subfamily transcription factors for creating high-performance biosensors. Notably, a similar insertion site was used for the LldR and PdhR, suggesting that it may be possible to create a wider range of biosensors using the same cpFP insertion site with other members of this protein family.

As with many other single FP-based biosensors, one of the limitations of the pyruvate biosensors is their pH sensitivity. The fluorescence intensities of GreenPy1 and ApplePy1 are largely dependent on the pH value (**Fig 2G,H**). In an effort to address this issue, we developed control biosensors, dGreenPy and dApplePy which have similar p*K*_a_ values to the unbound state of the parent biosensors and do not respond to changes in pyruvate concentration. Parallel imaging using these control biosensors can be used to identify artifacts caused by pH fluctuations. Another limitation is selectivity. GreenPy1 and ApplePy1 especially exhibit non-negligible responses to lactate, with the sensitivity to lactate increasing in proportion to their sensitivity to pyruvate. However, even the highest-affinity variants have high apparent *K*_d_s for L-lactate (∼ 10 mM range), and the lowest-affinity variants have even higher apparent *K*_d_s for L-lactate (> 100 mM). By selecting the appropriate affinity variant, interference from L-lactate can probably be avoided for most imaging experiments under typical physiological conditions.

In conclusion, we have developed the green fluorescent GreenPy1 series and red fluorescent ApplePy1 series of high-performance pyruvate biosensors. As the latest additions to the growing toolbox of FP-based metabolite biosensors, which also includes biosensors for L-lactate^9,10^, citrate^8^, and a variety of other metabolites^6,7^, these new tools will enable new insights and discoveries by providing researchers with improved capabilities to investigate metabolic process.

## Methods

### General methods and materials

A synthetic human codon-optimized gene encoding the *E. coli* PdhR transcriptional regulator protein was purchased from Integrated DNA Technologies. Phusion high fidelity DNA polymerase (Thermo Fisher Scientific) was used for routine polymerase chain reaction (PCR) amplification, and Taq DNA polymerase (New England Biolabs) was used for error-prone PCR. QuickChange mutagenesis kit (Agilent Technologies) was used for site-directed mutagenesis. Restriction endonucleases, rapid DNA ligation kits and GeneJET miniprep kits (Thermo Fisher Scientific) were used for plasmid construction and purification. Products of PCR and restriction digests were purified using agarose gel electrophoresis and the GeneJET gel extraction kit (Thermo Fisher Scientific). DNA sequences were analyzed by DNA sequence service of Fasmac Co. Ltd. Fluorescence spectra and intensity were recorded on Spark plate reader (Tecan). Genes encoding PyronicSF and Green Pegassos were obtained from Addgene (Addgene plasmid #124812 and #163114, respectively).

### Structural modeling

The modeling structures were generated by predicted by AlphaFold3^18^ by using AlphaFold3 server. The structure was analyzed using PyMol v.3.0.3 software.

### Engineering of GreenPy and ApplePy variants

The gene encoding cpGFP with N- and C-terminal linkers (DWS and NNV, respectively) was amplified using iGluSnFR gene as a template, followed by insertion into the indicated site of PdhR-PBD in a pBAD vector (Life Technologies) by Gibson assembly (New England Biolabs). The DNA binding domain of PdhR was removed beforehand. Variants were expressed in *E. coli* strain DH10B (Thermo Fisher Scientific) in LB media supplemented with 100 µg mL^-1^ ampicillin and 0.02% L-arabinose. Proteins were extracted using B-PER (bacterial protein extraction reagent, Thermo Fisher Scientific) for the assay of fluorescence brightness and pyruvate-dependent response. The most promising variant, designated as GreenPy0.1, was subjected to an iterative process of library generation and screening in *E. coli*. Libraries were generated by site-saturation mutagenesis^33^ to fully randomize the positions using an NNK codon or error-prone PCR of the whole gene with MnCl_2_ concentration adjusted to obtain one to two mutations per 1,000 base pairs. Primary screening was done on the agar plates, where approximately 2 × 10^3^ colonies were visually inspected each round. Total 384 bright colonies were then picked up and the bacteria were grown in 96-deepwell plates with 1 mL of LB supplemented with 100 μg mL^-1^ ampicillin and 0.02% L-arabinose at 37°C overnight, then transferred to room temperature and incubated overnight. Proteins were extracted with B-PER and the fluorescence brightness and the pyruvate-dependent response was assayed using a plate reader (Tecan). From this high-throughput assay, around 15 promising variants were selected and cultured again in 4 mL of LB. Proteins were extracted with B-PER and purified by Ni-NTA affinity chromatography (Agarose Bead Technologies). The brightness and the response were assayed again to select the template for the next library generation.

To create a gene encoding ApplePy0.1, the gene encoding cpRFP with N- and C-terminal linkers (DW and EATR, respectively) was amplified using R-iLACCO2 gene as a template, followed by insertion into the target site of PdhR-LBD in a pBAD vector by Gibson assembly. For linker optimization of ApplePy0.1, a variant with deleted N-linker was created by site-directed mutagenesis. In addition to it, a library where one residue linker was encoded using NNK codon was generated by QuickChange mutagenesis kit. The PCR product was used to transform *E*.*coli* and 96 brightly red fluorescent colonies were picked and cultured with 1 mL of LB supplemented with 100 μg mL^-1^ ampicillin and 0.02% L-arabinose at 37°C overnight, then transferred to room temperature and incubated overnight. Proteins were extracted with B-PER and the fluorescence brightness and the pyruvate-dependent response was assayed using a plate reader (Tecan). In parallel, libraries with two NNK codons and three NNK codons as N-linkers were also generated and screened similarly from 384 colonies. Proteins of promising variants selected from these libraries were extracted with B-PER and the brightness and the pyruvate-dependent response were assayed again to identify ApplePy0.2. To optimize the C-linker, two residues were randomized using NNK codons and 384 colonies were picked and screened in the same way. Following the linker optimization, directed evolution was performed by generating libraries by error-prone PCR and screening them in the same way as linker optimization of ApplePy. Subsequently, directed evolution was performed again from the 4th round variant. Libraries were generated by site-directed mutagenesis, site-saturation mutagenesis and random mutagenesis and screened as described above in the engineering of GreenPy variants.

To develop affinity variants and control biosensors, the mutations were introduced by site-directed mutagenesis and the response was assayed with purified proteins as described above, or libraries were generated by random-mutagenesis of the whole gene and screened in the same way under low pyruvate concentration (100 µM ∼ 300 µM).

### Protein purification and *in vitro* characterization

Pyruvate biosensors in pBAD expression vectors containing an N-terminal poly His (6×) tag were expressed in *E. coli* DH10B (Thermo Fisher Scientific). The bacteria were cultured in LB media supplemented with 100□μg□mL^−1^ ampicillin at 37 °C until the OD600 value of 0.6 was reached. The cell culture was then induced by adding 0.02% L-arabinose and grown overnight. Cell pellets were lysed with a cell disruptor (Branson), and proteins were purified by Ni-NTA affinity chromatography (Agarose Bead Technologies). Following purification the buffer was exchanged to 300□mM MOPS (pH 7.2) with 100□mM KCl. For pyruvate titration, buffers were prepared by mixing pyruvate (−) buffer (30□mM MOPS, 100□mM KCl, pH 7.2) and pyruvate (+) buffer (30□mM MOPS, 100□mM KCl, 100□mM pyruvate, pH 7.2) to provide pyruvate concentrations ranging from 0□mM to 100□mM. The excitation, emission and absorption spectra were all obtained using this buffer composition. To perform pH titrations, 10 µL protein solutions were diluted into 90 µL buffers that contained 30□mM trisodium citrate, 30□mM sodium borate, 30□mM MOPS, 100□mM KCl and either no pyruvate or 100□mM sodium pyruvate, and that had been adjusted to pH values ranging from 3 to 13. Absorbance spectra were measured using the Shimadzu UV1800 spectrometer and molar effective extinction coefficients were determined as previously reported^34^. Fluorescence quantum yields were measured using a Hamamatsu Photonics absolute quantum yield spectrometer (C9920-02G) using an excitation wavelength of 480 nm for green biosensors and 560 nm for red ones.

### Characterization of two-photon absorption properties

To measure molecular two-photon absorption cross sections of the chromophore in its deprotonated, anionic, state or protonated, neutral, state we need to know the molar concentration of these states. Therefore, here we define ε_A_ and ε_N_ as molar extinctions of either anionic or neutral forms of chromophore. The peak extinction coefficients of the anionic and neutral states were obtained using gradual alkaline titration method that measures the change of absorption spectrum as a function of pH^35^. The fractional concentrations of the anionic and neutral states, ρ_A_ and ρ_N_, respectively, were found using the Beer’s law from the extinction coefficients in both states and the total chromophore concentration. Absorption spectra were measured with a Lambda900 spectrophotometer (Perkin Elmer). The effective extinction coefficient defined above in the ***In vitro* characterization of GreenPy and ApplePy variants** section is calculated per molar concentration of total chromophore (both in anionic and neutral states) and, therefore, corresponds to a product ε_eff,A_ = ε_A_ ρ_A_ (for the “anionic” absorption peak) and ε_eff,N_ = ε_N_ρ_N_ (for the “neutral” absorption peak).

The methods and protocols of the two-photon characterization were described earlier^36^. Briefly, the two-photon absorption spectra were measured as corrected fluorescence excitation spectra by stepwise tuning the wavelength of a femtosecond laser (DeepSee Insight, MKS - Spectra-Physics) using a LabView program and collecting fluorescence signal at each wavelength with a PC1 spectrofluorimeter (ISS). The power dependence of fluorescence signal was checked for several excitation wavelengths and only the data points where it was quadratic were presented. For spectral shape measurements, a combination of short pass filters (770SP and 680SP) was used to block the scattered laser light. The cross-section σ_2,A_ of the anionic form was measured as described previously^37^, using rhodamine 6G in methanol as a reference standard, either 940 nm or 1060 nm for the green and red biosensors, respectively. At these wavelengths, only the anionic form gets excited. Molecular two-photon brightness evaluated at 940 (green biosensors) or 1060 nm (red biosensors) was calculated as a product of the anionic state fractional concentration ρ_A,_ fluorescence quantum yield φ_A_, and two-photon cross section σ_2,A_: F_2,A_ = φ_A_ σ_2,A_ ρ_A_. To obtain the two-photon excitation spectra in units of molecular brightness, the corrected (for the spectral variations of laser properties), unscaled two-photon excitation spectra were scaled to the product F_2,A_ values, measured at 940 nm (green biosensors) or 1060 nm (red biosensors). To directly measure the signal-to background ratio upon saturating with pyruvate, the fluorescence signals (either under one-photon or two-photon excitation) were measured in the pyruvate-free and pyruvate-saturated samples with the same concentration of protein and under the same conditions.

### Construction of mammalian expression vectors

The genes of GreenPy variants, ApplePy variants, the gene of PyronicSF (Addgene plasmid no. 124812), and the gene of and Green Pegassos (Addgene plasmid no. 163114) were ligated into pcDNA3 vector (Thermo Fisher Scientific) with T4 ligase (Thermo Fisher Scientific). To construct the plasmids for expression in mitochondria, two-times repeated COX8 sequences were fused at the N-terminus of the gene of interest.

### Imaging of the GreenPy1 series and ApplePy1 series in HeLa cells

HeLa (American Type Culture Collection; ATCC #CCL-2) cells were maintained in Dulbecco’s modified Eagle medium (DMEM high glucose; Nacalai Tesque) supplemented with 10% fetal bovine serum (FBS; Sigma-Aldrich), and 100 μg mL^-1^ penicillin and streptomycin (Nacalai Tesque). Cells were transiently transfected with the plasmids with Lipofectamine® 3000 Reagent (Thermo Fisher Science) in Opti-MEM (Gibco) and imaged within 24 - 72 hours after transfection. IX83 wide-field fluorescence microscopy (Olympus) equipped with a pE-300 LED light source (CoolLED), and a 40× objective lens (numerical aperture (NA) = 1.3; oil), an ImagEM X2 EMCCD camera (Hamamatsu), and Cellsens software (Olympus) were used for the imaging. The filter sets in the imaging were the following specification. ApplePy: excitation 545/20 nm, dichroic mirror 565 nm dclp, and emission 598/55 nm; GreenPy variants, PyronicSF, and Green Pegassos: excitation 470/20 nm, dichroic mirror 490 nm dclp, and emission 518/45 nm. Fluorescence images were analyzed with ImageJ software (https://imagej.net/software/fiji/, National Institutes of Health).

For imaging in treatment with 10 mM sodium pyruvate, MCT inhibitor AR-C155858 (Tocris) and MPC inhibitor UK 5099 (MedChemExpress), Hank’s balanced salt solution (HBSS; without phenol red, Nacalai Tesque) supplemented with 10 mM HEPES (Nacalai Tesque) was used as an imaging buffer. The reagents of UK 5099 and AR-C155858 were stored in the concentration of 10 mM in DMSO. The final concentration of UK 5099 and AR-C155858 in HeLa cells are 25 μM and 1 μM respectively. For the removal of intracellular pyruvate, HeLa cells were preincubated with 500 μM iodoacetic acid in DMEM for 15 - 30 mins, and the imaging was carried out in the imaging buffer contained 10 μM nigericin and 2 μM rotenone. In starvation experiments, HeLa cells were incubated in no-glucose DMEM (Nacalai Tesque) for 2 - 3 hours before imaging. After exchanging medium into no-glucose imaging buffer (184.45 mg L^-1^ CaCl_2_·H_2_O, 97.6 mg L^-1^ MgSO_4_, 400.00 mg L^-1^ KCl, 60.00 mg L^-1^ KH_2_PO_4_, 8000.00 mg L^-1^ NaCl, 350.00 mg L^-1^ NaHCO_3_, 47.88 mg L^-1^ Na_2_HPO_4_), final concentration of 5 mM D-glucose was added at time = 0 min. In all the imaging, snapshots were taken every five seconds. All the reagents were diluted up to the ten times concentration of required final concentration, and 200 μL of solution was added to 1.8 mL of imaging buffer. All the cell imaging was repeated at least 3 times independently and the number of cells that we analyzed were shown on figure legend.

## Data availability

The data that support the findings of this study are available from the corresponding authors on reasonable request. The bacterial and mammalian cell expression plasmids encoding GreenPy1 and ApplePy1 variants are available from Addgene (#234049 - #234078; https://www.addgene.org/browse/article/28253502/).

## Author contributions

S.I. and C.S. developed GreenPy variants. S.I., I.Y., and K.S. developed ApplePy variants.

S.I. performed *in vitro* characterization. S.H. performed cell imaging and data analysis. M.D.,

A.T and B.M. characterized two-photon absorption properties (spectra and molecular brightness). K.T-Y., T.T., and R.E.C. supervised research. S.I., S.H., and R.E.C. wrote the manuscript.

## Acknowledgements

This research was supported by the Japan Society for the Promotion of Science (JSPS) (24KJ0642 to S.H., 21H00273 and 23H02101 to T.T., and 19H05633, 22H04743, 24H00489, and 24H02267 to R.E.C.). S.I. was supported by the MERIT-WINGS program of The University of Tokyo and S.H. was supported by JST SPRING (JPMJSP2108). Resource for Multiphoton Characterization of Genetically Encoded Probes at Montana State University was supported by the NIH/NINDS grant U24NS109107 under the BRAIN Initiative.

## References

1. Gray, L. R., Tompkins, S. C. & Taylor, E. B. Regulation of pyruvate metabolism and human disease. Cell. Mol. Life Sci. 71, 2577–2604 (2014).

2. Gest, A. M. M. et al. Molecular spies in action: Genetically encoded fluorescent biosensors light up cellular signals. Chem. Rev. 124, 12573–12660 (2024).

3. Nasu, Y., Shen, Y., Kramer, L. & Campbell, R. E. Structure- and mechanism-guided design of single fluorescent protein-based biosensors. Nat. Chem. Biol. 17, 509–518 (2021).

4. Zhang, Y. et al. Fast and sensitive GCaMP calcium indicators for imaging neural populations. Nature 615, 884–891 (2023).

5. Feng, J. et al. Monitoring norepinephrine release in vivo using next-generation GRABNE sensors. Neuron 112, 1930-1942.e6 (2024).

6. Zhang, Z., Cheng, X., Zhao, Y. & Yang, Y. Lighting Up Live-Cell and In Vivo Central Carbon Metabolism with Genetically Encoded Fluorescent Sensors. Annu. Rev. Anal. Chem. 13, 293–314 (2020).

7. Díaz-García, C. M. et al. Quantitative in vivo imaging of neuronal glucose concentrations with a genetically encoded fluorescence lifetime sensor. J. Neurosci. Res. 97, 946–960 (2019).

8. Zhao, Y., Shen, Y., Wen, Y. & Campbell, R. E. High-Performance Intensiometric Direct- and Inverse-Response Genetically Encoded Biosensors for Citrate. ACS Cent Sci 6, 1441–1450 (2020).

9. Hario, S. et al. High-performance genetically encoded green fluorescent biosensors for intracellular l-lactate. ACS Cent. Sci. 10, 402–416 (2024).

10. Nasu, Y. et al. Lactate biosensors for spectrally and spatially multiplexed fluorescence imaging. Nat. Commun. 14, 6598 (2023).

11. Bulusu, V. et al. Spatiotemporal analysis of a glycolytic activity gradient linked to mouse embryo mesoderm development. Dev. Cell 40, 331-341.e4 (2017).

12. Peroza, E. A.Boumezbeur, A.-H. & Zamboni, N. Rapid, randomized development of genetically encoded FRET sensors for small molecules. Analyst 140, 4540–4548 (2015).

13. San Martín, A. et al. Imaging mitochondrial flux in single cells with a FRET sensor for pyruvate. PLoS One 9, e85780 (2014).

14. Arce-Molina, R. et al. A highly responsive pyruvate sensor reveals pathway-regulatory role of the mitochondrial pyruvate carrier MPC. Elife 9, (2020).

15. Harada, K. et al. Green fluorescent protein-based lactate and pyruvate indicators suitable for biochemical assays and live cell imaging. Sci. Rep. 10, 19562 (2020).

16. Quail, M. A. & Guest, J. R. Purification, characterization and mode of action of PdhR, the transcriptional repressor of the pdhR?aceEF?Ipd operon of Escherichia coli. Mol. Microbiol. 15, 519–529 (1995).

17. Suvorova, I. A., Korostelev, Y. D. & Gelfand, M. S. GntR family of bacterial transcription factors and their DNA binding motifs: Structure, positioning and co-evolution. PLoS One 10, e0132618 (2015).

18. Abramson, J. et al. Accurate structure prediction of biomolecular interactions with AlphaFold 3. Nature 630, 493–500 (2024).

19. Marvin, J. S. et al. An optimized fluorescent probe for visualizing glutamate neurotransmission. Nat. Methods 10, 162–170 (2013).

20. Zhu, A., Romero, R. & Petty, H. R. A sensitive fluorimetric assay for pyruvate. Anal. Biochem. 396, 146–151 (2010).

21. athioudakis, D. et al. Pyruvate: immunonutritional effects on neutrophil intracellular amino or alpha-keto acid profiles and reactive oxygen species production. Amino Acids 40, 1077–1090 (2011).

22. Parnetti, L. et al. Increased cerebrospinal fluid pyruvate levels in Alzheimer’s disease. Neurosci. Lett. 199, 231–233 (1995).

23. Ahmed, S. S., Santosh, W., Kumar, S. & Christlet, H. T. T. Metabolic profiling of Parkinson’s disease: evidence of biomarker from gene expression analysis and rapid neural network detection. J. Biomed. Sci. 16, 63 (2009).

24. Clarke, C. et al. Mitochondrial respiratory chain disease discrimination by retrospective cohort analysis of blood metabolites. Mol. Genet. Metab. 110, 145–152 (2013).

25. Kalivoda, K. A., Steenbergen, S. M. & Vimr, E. R. Control of the Escherichia coli sialoregulon by transcriptional repressor NanR. J. Bacteriol. 195, 4689–4701 (2013).

26. San Martín, A. et al. A genetically encoded FRET lactate sensor and its use to detect the Warburg effect in single cancer cells. PLoS One 8, e57712 (2013).

27. Poole, R. C. & Halestrap, A. P. Transport of lactate and other monocarboxylates across mammalian plasma membranes. Am. J. Physiol. 264, C761–82 (1993).

28. Keller, J. P. et al. In vivo glucose imaging in multiple model organisms with an engineered single-wavelength sensor. Cell Rep. 35, 109284 (2021).

29. Marvin, J. S. et al. A genetically encoded fluorescent sensor for in vivo imaging of GABA. Nat. Methods 16, 763–770 (2019).

30. Sun, F. et al. A genetically encoded fluorescent sensor enables rapid and specific detection of dopamine in flies, fish, and mice. Cell 174, 481-496.e19 (2018).

31. Patriarchi, T. et al. Ultrafast neuronal imaging of dopamine dynamics with designed genetically encoded sensors. Science 360, (2018).

32. Kubitschke, M. et al. Next generation genetically encoded fluorescent sensors for serotonin. Nat. Commun. 13, 7525 (2022).

33. Ho, S. N., Hunt, H. D., Horton, R. M., Pullen, J. K. & Pease, L. R. Site-directed mutagenesis by overlap extension using the polymerase chain reaction. Gene 77, 51– 59 (1989).

34. Cranfill, P. J. et al. Quantitative assessment of fluorescent proteins. Nat. Methods 13, 557–562 (2016).

35. Barnett, L. M., Hughes, T. E. & Drobizhev, M. Deciphering the molecular mechanism responsible for GCaMP6m’s Ca2+-dependent change in fluorescence. PLoS One 12, e0170934 (2017).

36. Drobizhev, M., Molina, R. S. & Hughes, T. E. Characterizing the Two-photon Absorption Properties of Fluorescent Molecules in the 680-1300 nm Spectral Range. Bio Protoc. 10, e3498 (2020).

37. Dalangin, R. et al. Far-red fluorescent genetically encoded calcium ion indicators. Nat. Commun. 16, 3318 (2025).

