## Supplementary Information for "High-performance genetically-encoded green and red fluorescent biosensors for pyruvate"

### Supplementary Notes

#### Note S1. Development of ApplePy0.9 via directed evolution

Starting from ApplePy0.3, we focused on improving the $\Delta F/F_0$. Initially, we conducted directed evolution by screening libraries of randomly mutated variants. Using crude lysate assays, we achieved a $\Delta F/F_0$ of 16.8 after 17 iterative rounds. The final variant carried the following mutations: F5S, E11G, L13F, D15Y, T43S, D46E, L23M, L90M, L93V, N103H, I132V, K139T, K153M, K169M, M197L, K202M, Y311C, K383R, R393K, and E408G. However, we later found that the $\Delta F/F_0$ for the purified protein was roughly half of that observed in crude lysate, and expression in HeLa cells was poor.

To assess whether a particularly detrimental mutation had been introduced during the 17 rounds of directed evolution, we performed detailed *in vitro* and mammalian cell-based characterization of the best variant from each round. This analysis revealed that variants from round 5 onward all exhibited the same issues as the final variant. Consequently, we restarted directed evolution using the round 4 variant, which included the mutations D46E, K169M, M197L, and E408G, and modified the protocol to assay purified proteins rather than crude lysates generated by detergent-mediated lysis.

To build the insights obtained from the previous 17 rounds of directed evolution, we began with site-saturation mutagenesis of residues L93 and N103, which had previously led to a significant increase in $\Delta F/F_0$. However, this effort did not lead to the identification of improved variants. We then performed four rounds of whole-gene directed evolution, yielding the mutations S193P and Q106H. Subsequent site-saturation mutagenesis targeting residues S42 and T43, which had seemed to potentially be beneficial based on the previous directed evolution, also failed to produce improved variants.

We then shifted focus to residues E357 and A358, located in the C-linker and had not been previously mutated during linker optimization, and identified the beneficial A358S mutation. Next, we used site-directed mutagenesis to test the effects of L13F, D15Y, T43S, L93V, I132V, K383E, and R393K. Among these, the combination of T43S and I132V yielded the most improved variants. Finally, we conducted two additional rounds of whole-gene directed evolution, introducing the mutations C95Y, Y227F, K351M, E399K, R414L, and K423E. This final variant was designated as ApplePy0.9.

#### Note S2. Affinity tuning of GreenPy and ApplePy biosensors

We first focused on developing affinity variants of ApplePy0.9, which had an apparent $K_d$ of ~ 700 μM. We initially introduced the S369A mutation, which was identified in the final round of directed evolution. The resulting variant exhibited lower affinity and a much higher $\Delta F/F_0$ and was designated ApplePy1Lowest. For all further work we focused on increasing the affinity of ApplePy1Lowest rather than ApplePy0.9.

During the development of iLACCO affinity variants[1], V100T, V100I, V393T, and V393I had been rationally selected as candidates to alter affinity without affecting the fluorescence change. Given the high similarity between LldR and PdhR, these residues were expected to impact affinity. Indeed, according to the predicted structure (**Supplementary Figure 9**), both residues 100 and 393 point toward the putative binding pocket. Therefore, we tested the corresponding mutations (V104I, V104T, I398V, and I398T) in ApplePy0.9, expecting to alter the affinity. As a result, we obtained a higher affinity variant with the V104I mutation, designated ApplePy1Low.

To further increase the affinity, we performed one round of directed evolution with error-prone PCR of the whole gene, and identified ApplePy1High, which carried the V10L mutation and exhibited similar $\Delta F/F_0$ with higher affinity. At the same time, we identified several additional mutations that increased affinity. These were introduced into ApplePy1High, and their performance was tested individually. Among them, ApplePy1High-E58G showed the highest affinity, though it exhibited a lower response. To recover the response of ApplePy1High-E58G, we performed two additional rounds directed evolution with error-prone PCR of the whole gene, and obtained ApplePy1Highest, which carried the L200Q, Y201F, and P344L mutations.

Next, we turned our attention to the affinity tuning of GreenPy0.9. Based on accumulated results from random mutagenesis of both ApplePy and GreenPy, we selected the following mutations as candidates to alter affinity for pyruvate: V10L, Q20H, L74F, I78T, I104V, Q106H, S366A, and H371N. Their effects were tested individually, and V10L, E58G, I78T, I104V, and S366A were selected for further investigation. We created single, double, or triple mutants by combining these mutations and assayed their affinity and fluorescence changes in detail. This led to the identification of GreenPy1Lowest (GreenPy0.9-E58G-I78T-I104V), GreenPy1Low (GreenPy0.9-I78T), and GreenPy1High (GreenPy0.9-E58G-I78T). We found that GreenPy0.9-E58G exhibited the highest affinity, though with a reduced response. Accordingly, we performed two additional rounds of directed evolution with error-prone PCR of the whole gene, and identified the variant that we designated as GreenPy1Highest, which carried the P17T mutation.

#### Supplementary Tables

##### Table S1. Detailed photophyisical properties of GreenPy1 variants

| Variant | GreenPy1 Highest | | GreenPy1 High | | GreenPy1 Low | | GreenPy1 Lowest | |
|---|---|---|---|---|---|---|---|---|
| Pyruvate | - | + | - | + | - | + | - | + |
| Relative fraction of neutral chromophore ($\rho_N$) | 0.92 | 0.41 | 0.98 | 0.40 | 0.97 | 0.44 | 0.97 | 0.51 |
| Relative fraction of anionic chromophore ($\rho_A$) | 0.08 | 0.59 | 0.024 | 0.60 | 0.03 | 0.56 | 0.03 | 0.49 |
| Neutral absoption peak (nm) | 385 | 387 | 385 | 385 | 385 | 385 | 385 | 385 |
| Anionic absoption peak (nm) | 482 | 483 | 501 | 484 | 500 | 483 | 500 | 483 |
| Neutral extinction coefficient ($\varepsilon_N$, $mM^{-1}cm^{-1}$) | 36 | 40 | 38 | 40 | 34 | 38 | 39 | 38 |
| Anionic extinction coefficient ($\varepsilon_A$, $mM^{-1}cm^{-1}$) | 49 | 60 | 60 | 67 | 64 | 59 | 61 | 73 |
| Anionic quantum yield ($\varphi_A$) | 0.46 | 0.52 | 0.30 | 0.48 | 0.33 | 0.49 | 0.35 | 0.52 |
| Anionic molecular brightness ($\rho_A$ $\varepsilon_A$ $\varphi_A$) | 1.80 | 18.4 | 0.43 | 19.3 | 0.63 | 16.2 | 0.64 | 18.6 |
| Anionic two-photon cross seciton (GM at λ nm) | 25 (940) | 30 (940) | 29 (940) | 34 (940) | 31 (940) | 32 (940) | 28 (940) | 35 (940) |
| Anionic two-photon brightness ($F_2$, GM at λ nm) | 0.92 (940) | 9.2 (940) | 0.21 (940) | 9.8 (940) | 0.31 (940) | 8.8 (940) | 0.29 (940) | 8.9 (940) |

**Table S2. Detailed photophyisical properties of ApplePy1 variants**

| Variant | ApplePy1 Highest | | ApplePy1 High | | ApplePy1 Low | | ApplePy1 Lowest | |
|---|---|---|---|---|---|---|---|---|
| Pyruvate | - | + | - | + | - | + | - | + |
| Relative fraction of neutral chromophore ($\rho_N$) | 0.92 | 0.63 | 0.97 | 0.58 | 0.97 | 0.60 | 0.96 | 0.70 |
| Relative fraction of anionic chromophore ($\rho_A$) | 0.08 | 0.37 | 0.03 | 0.42 | 0.03 | 0.40 | 0.04 | 0.30 |
| Neutral absoption peak (nm) | 446 | 471 | 446 | 472 | 445 | 455 | 444 | 455 |
| Anionic absoption peak (nm) | 556 | 554 | 567 | 554 | 569 | 554 | 571 | 555 |
| Neutral extinction coefficient ($\varepsilon_N$, $mM^{-1}cm^{-1}$) | 34 | 29 | 39 | 35.5 | 32.5 | 31.5 | 25 | 26 |
| Anionic extinction coefficient ($\varepsilon_A$, $mM^{-1}cm^{-1}$) | 94 | 88 | 92 | 95 | 91 | 86 | 68 | 76 |
| Anionic quantum yield ($\varphi_A$) | 0.23 | 0.33 | 0.15 | 0.34 | 0.16 | 0.36 | 0.12 | 0.35 |
| Anionic molecular brightness ($\rho_A$ $\varepsilon_A$ $\varphi_A$) | 1.73 | 10.7 | 0.41 | 14 | 0.44 | 12.3 | 0.33 | 8.0 |
| Anionic two-photon cross seciton (GM at λ nm) | 53 (1060) | 64 (1060) | 38 (1060) | 64 (1060) | 50 (1060) | 48 (1060) | 32 (1060) | 51 (1060) |
| Anionic two-photon brightness ($F_2$, GM at λ nm) | 0.98 (1060) | 7.8 (1060) | 0.17 (1060) | 9.1 (1060) | 0.24 (1060) | 6.9 (1060) | 0.15 (1060) | 5.4 (1060) |

#### Supplementary Figures

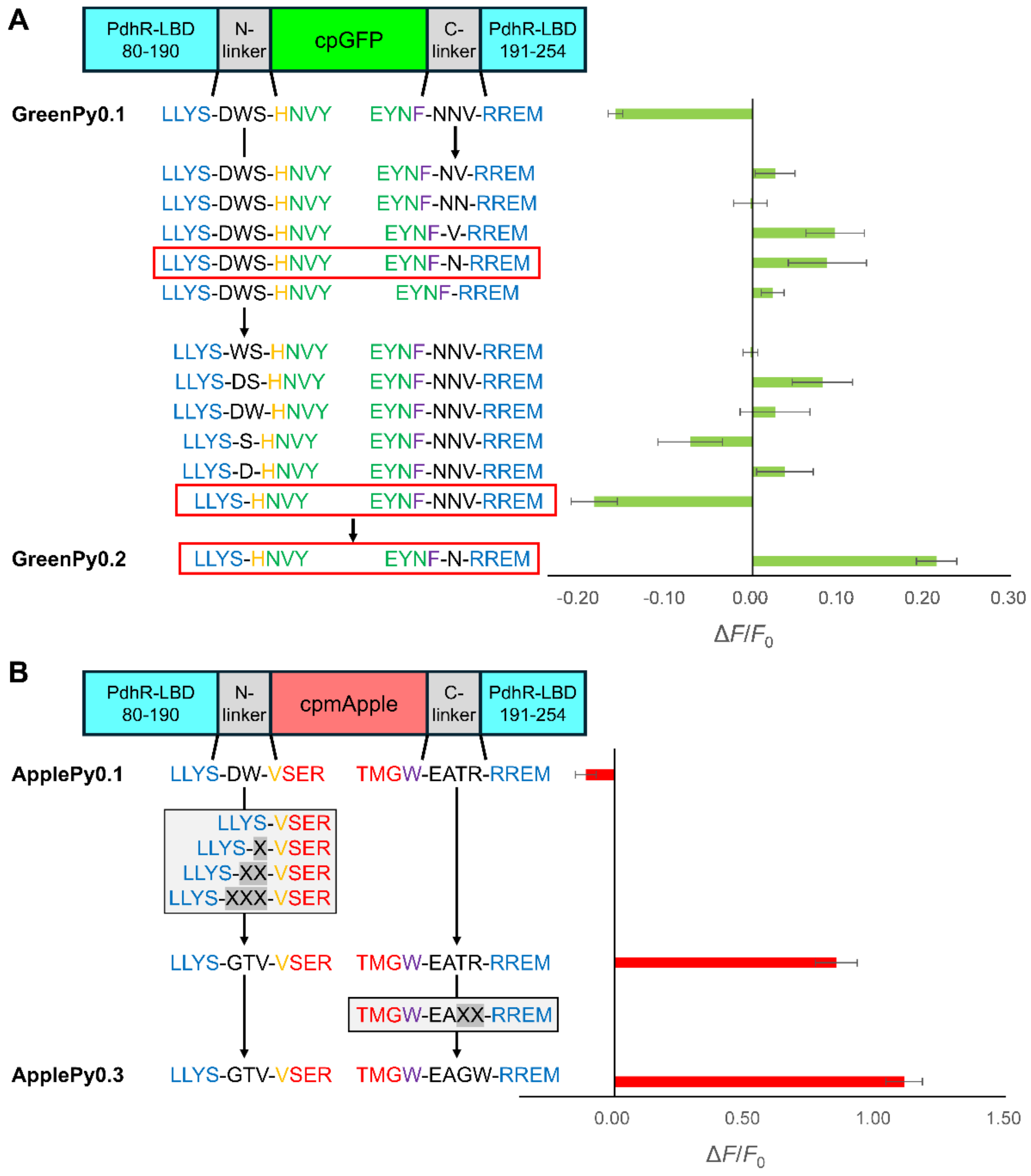

**Figure S1. Linker optimization**

**A,** $\Delta F/F_0$ of GreenPy variants with different linker lengths (n = 3 replicates; mean ± s.d.). When shortening the C-linker, two types of partially deleted variants (C-linker = V, N) exhibited the largest positive response, though the latter had higher brightness. When shortening the N-linker, total deletion yielded the largest fluorescence change. When these two were combined, the response increased to 0.21, and therefore this variant was designated as GreenPy0.2 for further improvement. **B,** Schematic representation of the strategy and results to optimize the linkers of ApplePy0.1. To optimize the N-linker, we optimized the length and linkers simultaneously by generating libraries in which the N-linker was encoded by 1 to 3 NNK degenerate codons. These three libraries were screened in parallel along with the no linker variant, and the most responsive variant was selected. Positions T359 and R360 were then subjected to site-saturation mutagenesis to optimize the sequence of the C-linker.

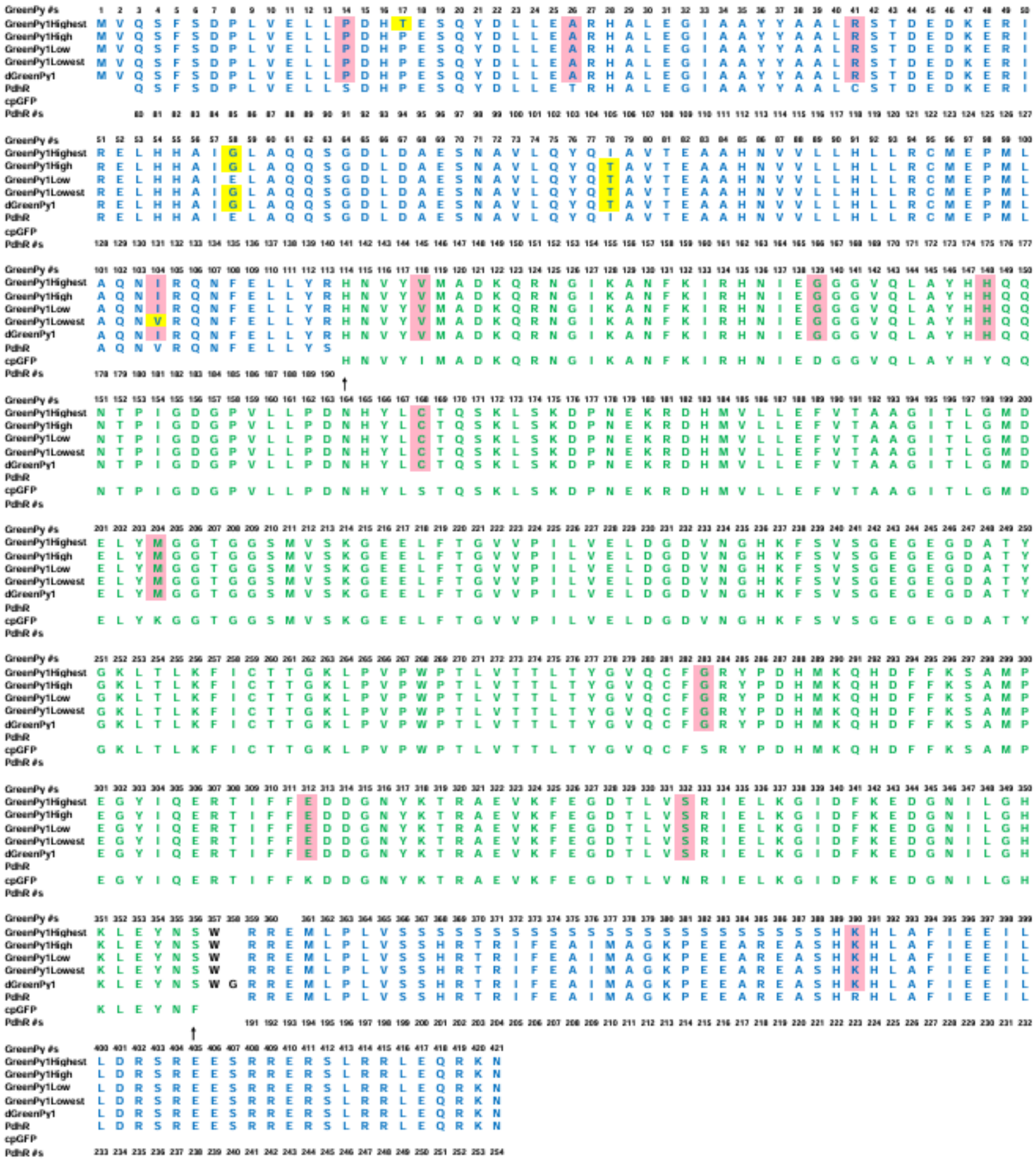

**Figure S2. Sequence alignment of GreenPy variants**

The PdhR-LBD domain is colored blue, cpGFP is colored green, and the linkers are colored black. Mutations in GreenPy0.9 relative to GreenPy0.2 are indicated by a red background, and mutations for affinity tuning are indicated by a yellow background. The gatepost residues are indicated by arrows.

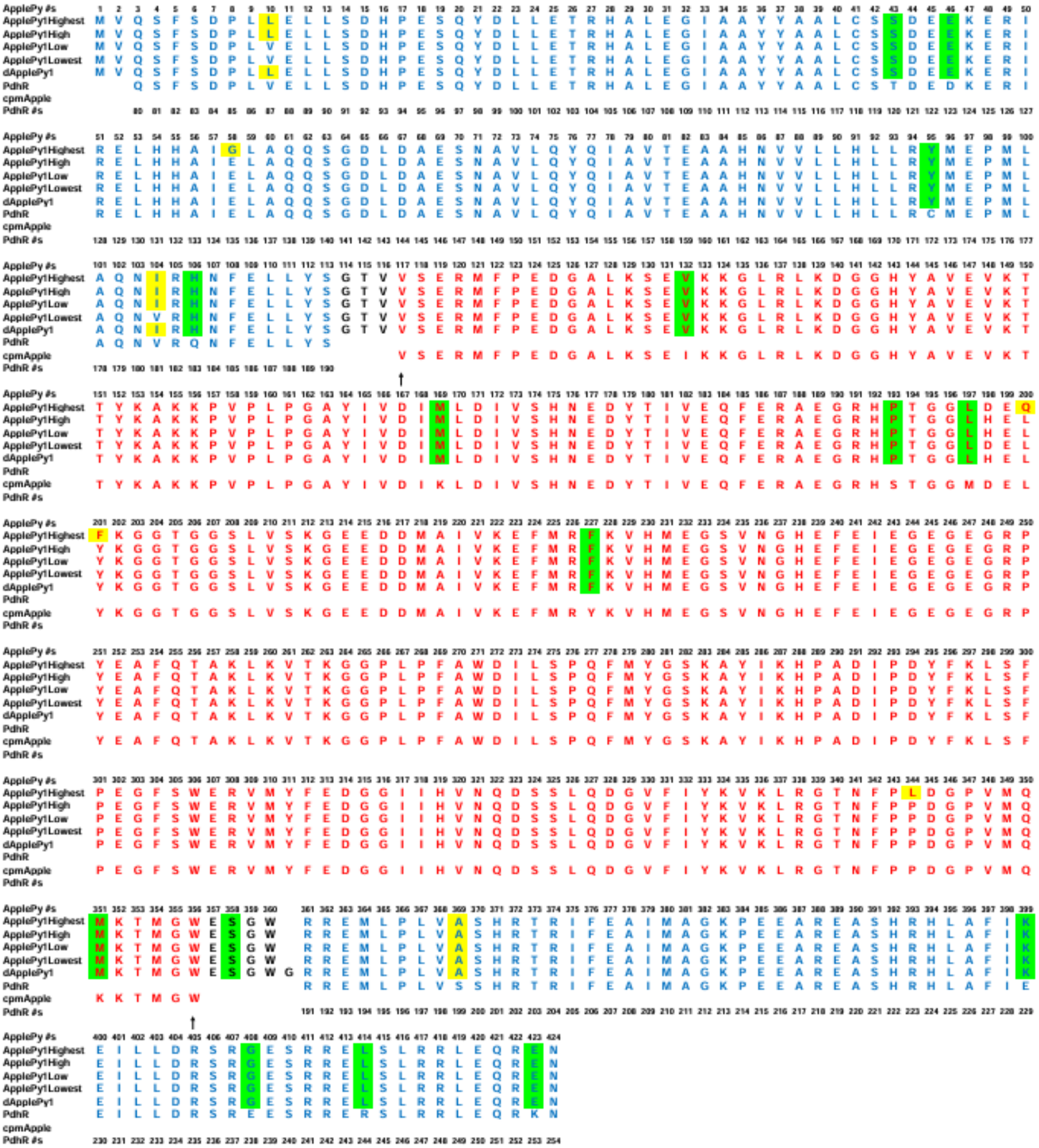

**Figure S3. Sequence alignment of ApplePy variants**

The PdhR-LBD is colored blue, cpmApple is colored red, and the linkers are colored in black. Mutations in ApplePy0.9 relative to ApplePy0.3 are indicated by a green background, and mutations for affinity tuning are indicated by a yellow background. The gatepost residues are indicated by arrows.

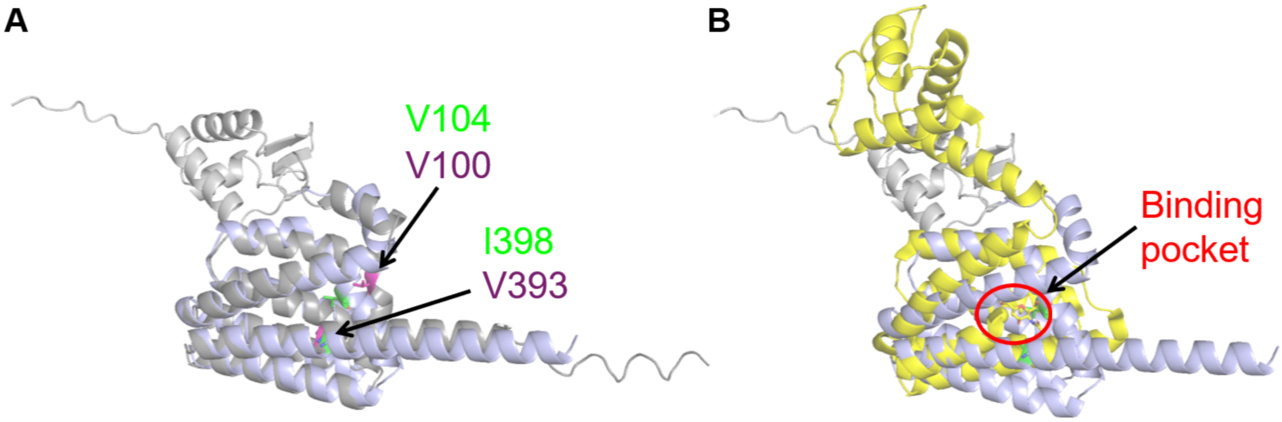

**Figure S4. Structure modeling for affinity tuning.**

**A,** The predicted structure of PdhR (LBD: blue, DBD: gray) superimposed with the predicted structure of LldR-LBD (gray). V100 and V393 of LldR (iLACCO1 numbering) are colored purple, and the corresponding V104 and I398 of PdhR (ApplePy1 numbering) are colored green. **B,** The predicted structure of PdhR (LBD: blue, DBD: gray) superimposed with a crystal structure of NanR (yellow, PDB ID: 6ON4)[2]. NanR is another transcription factor that belongs to the FadR subfamily and can bind to sialic acid[3,4]. The red circle indicates the binding pocket of NanR. V104 and I398 residues are located in this putative binding pocket.

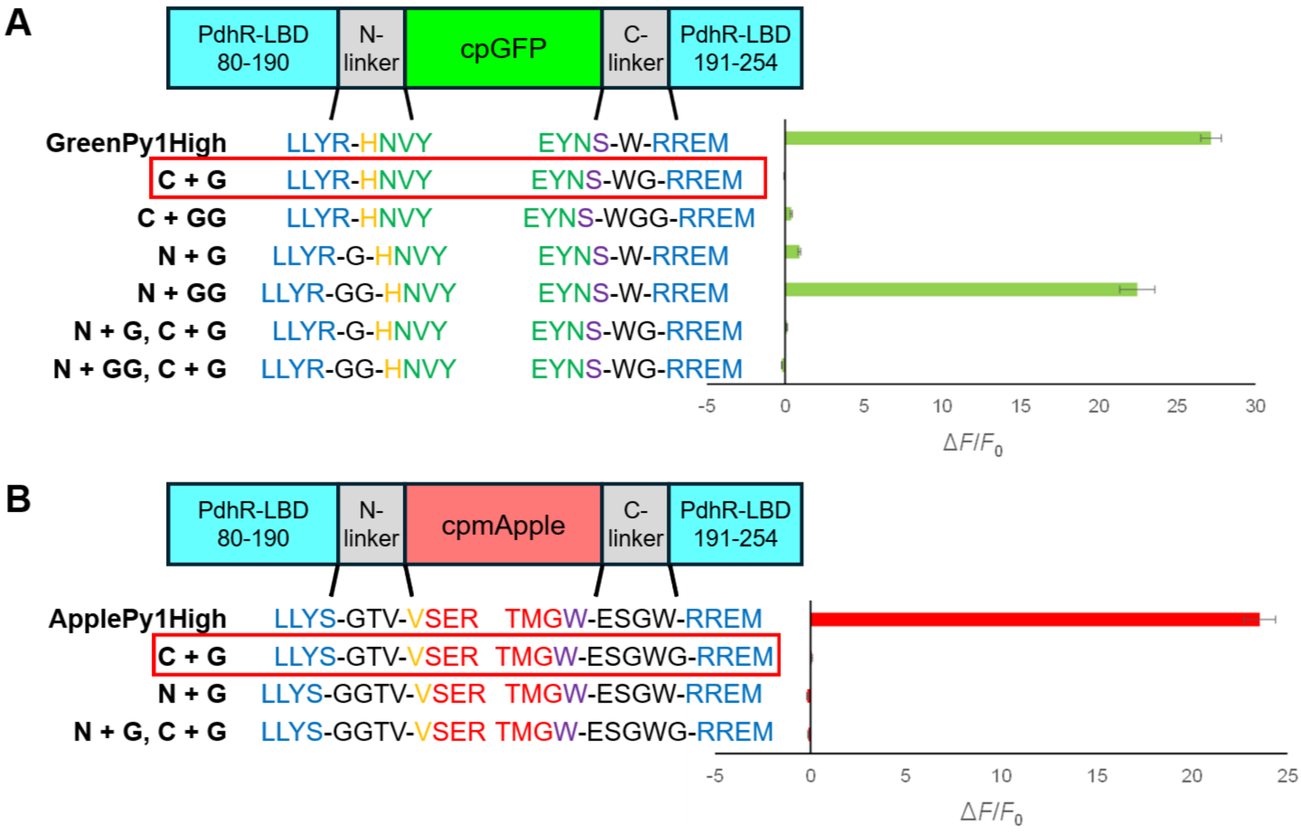

**Figure S5. Development of control biosensors**

**A,B,** Schematic representation of the strategy to develop control biosensors for the GreenPy1 series (**A**) and the ApplePy1 series (**B**). One or two glycine residues were inserted into the PdhR side of either or both linkers connecting to the PdhR domain. The linker length is generally critical for the function of biosensors, so it was expected that the change in the linker length might abolish the response while retaining the binding to pyruvate and the microenvironment around the chromophore. $\Delta F/F_0$ of variants with different linker length are shown in the bar graph (n = 3 replicates; mean ± s.d.). GreenPy1High-W357_R358insG (C+G) and ApplePy1High-W360_R361insG (C+G) exhibited the smallest response and similar brightness in the pyruvate free state and were designated as dGreenPy and dApplePy, respectively.

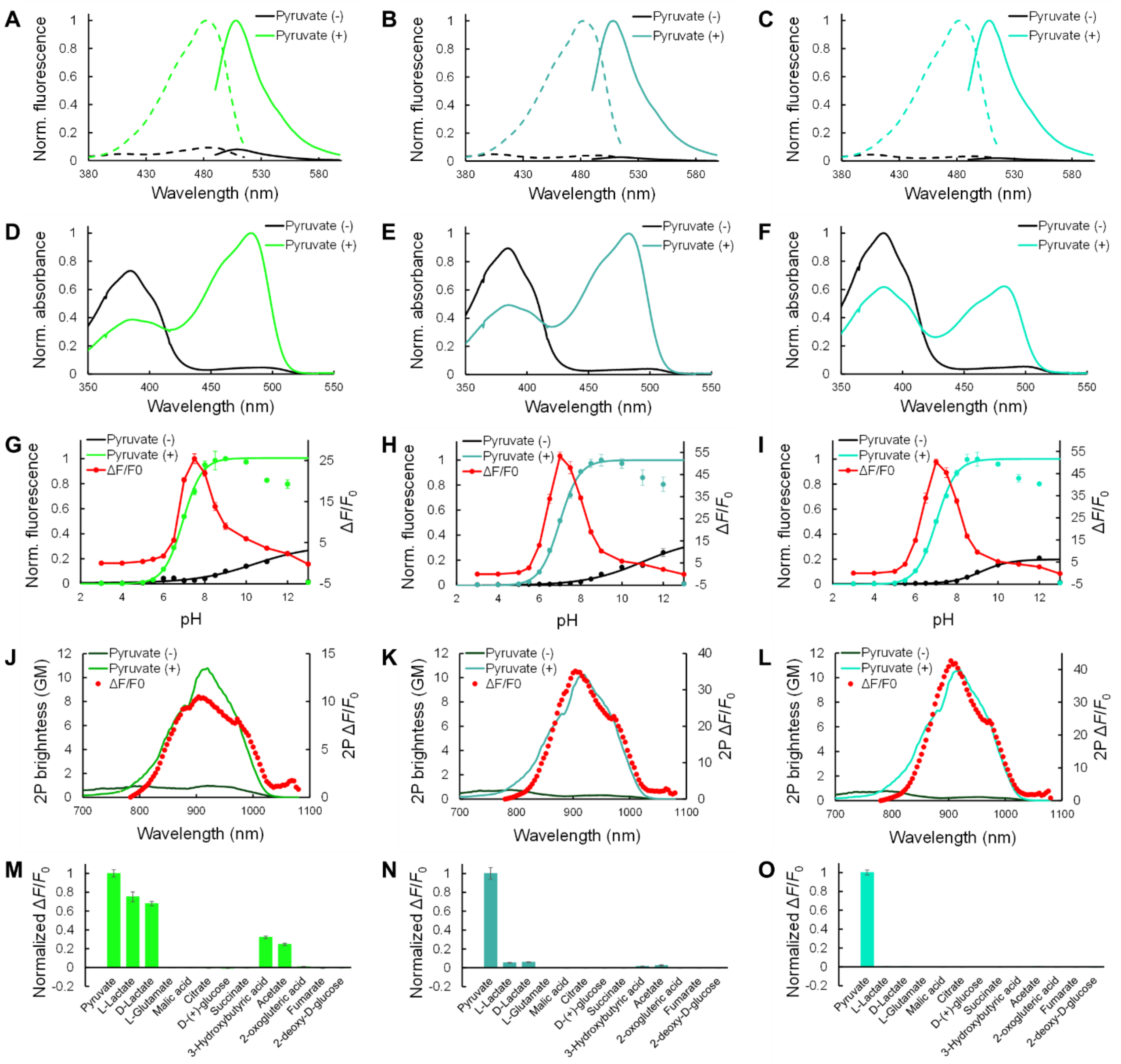

**Figure S6.** ***In vitro*** **characterization of GreenPy affinity variants**

**A-C,** Excitation (emission at 555nm) and emission (excitation at 450 nm) spectra of GreenPy1Highest (**A**), GreenPy1Low (**B**) and GreenPy1Lowest (**C**) in the presence (100 mM) and absence of pyruvate. **D-F,** Absorbance spectra of GreenPy1Highest (**D**), GreenPy1Low (**E**) and GreenPy1Lowest (**F**) in the presence (10 mM) and absence of pyruvate. **G-I,** pH titration curve of GreenPy1Highest (**G**), GreenPy1Low (**H**) and GreenPy1Lowest (**I**) in the presence (100 mM) and the absence of pyruvate (n = 3 replicates; mean ± s.d.). The curve was fitted to calculate the physiological p$K_a$ value. **J-L,** Two-photon excitation spectra of GreenPy1Highest (**J**), GreenPy1Low (**K**) and GreenPy1Lowest (**L**) in the presence (10 mM) and absence of pyruvate. 2P, Two-photon. GM, Goeppert-Mayer units. **M-O,** Fluorescence responses of GreenPy1Highest (**M**), GreenPy1Low (**N**) and GreenPy1Lowest (**O**) to various metabolites at 10 mM (n = 3 replicates; mean ± s.d.).

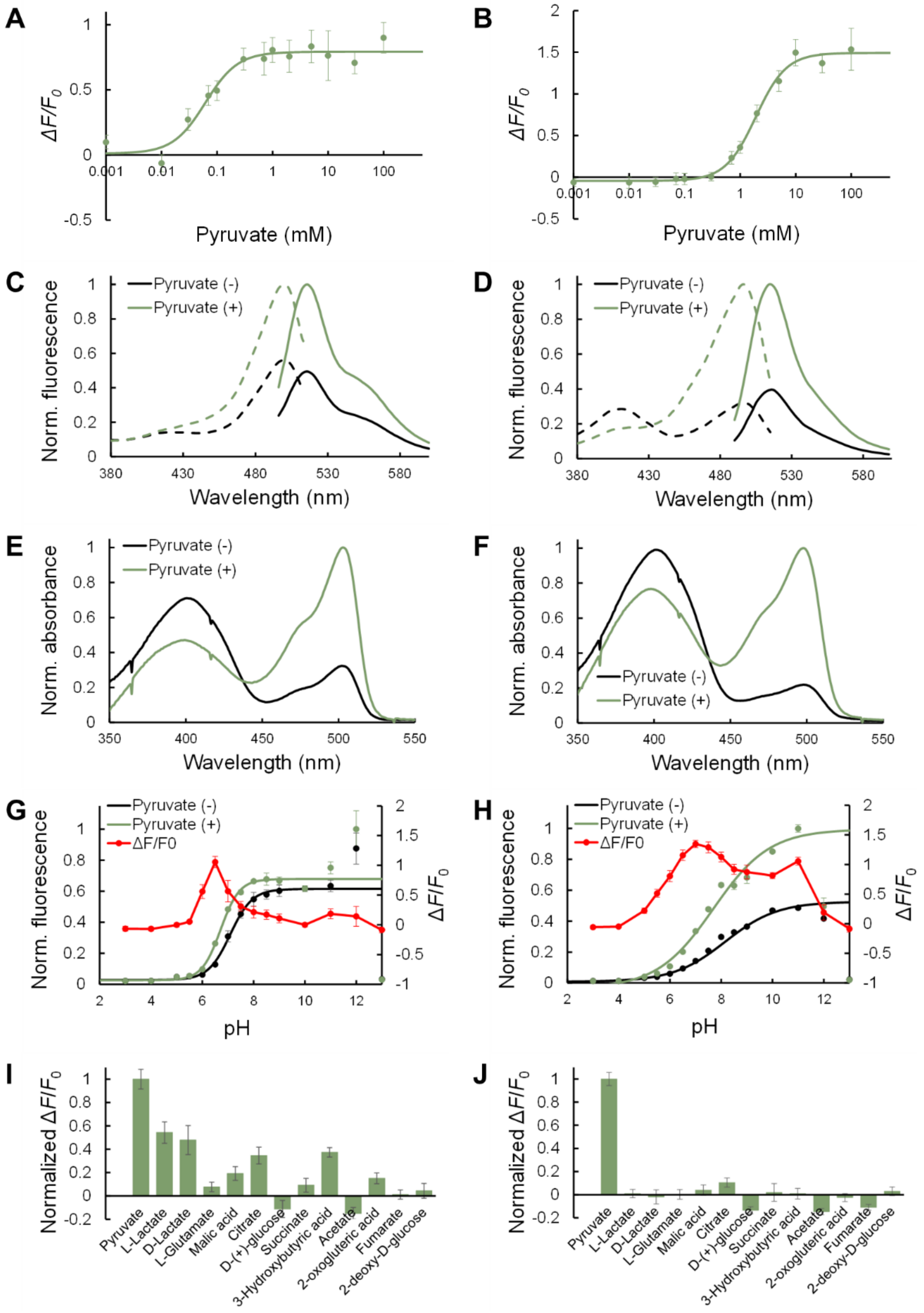

**Figure S7.** ***In vitro*** **characterization of previously reported green pyruvate biosensors**

**A,B,** Pyruvate titration curve of purified Green Pegassos (**A**) and PyronicSF (**B**) variants (n = 3 replicates; mean ± s.d.). **C,D,** Excitation (emission at 555 nm) and emission (excitation at 450 nm) spectra of Green Pegassos (**C**) and PyronicSF (**D**) in the presence (100 mM) and absence of pyruvate. **E,F,** Absorbance spectra of Green Pegassos (**E**) and PyronicSF (**F**) in the presence (10 mM) and absence of pyruvate. **G,H,** pH titration curve of Green Pegassos (**G**) and PyronicSF (**H**) in the presence (100 mM) and the absence of pyruvate (n = 3 replicates; mean ± s.d.). The curve was fitted to calculate the physiological p$K_a$ value. **I,J,** Fluorescence responses of Green Pegassos (**I**) and PyronicSF (**J**) to various metabolites (n = 3 replicates; mean ± s.d.).

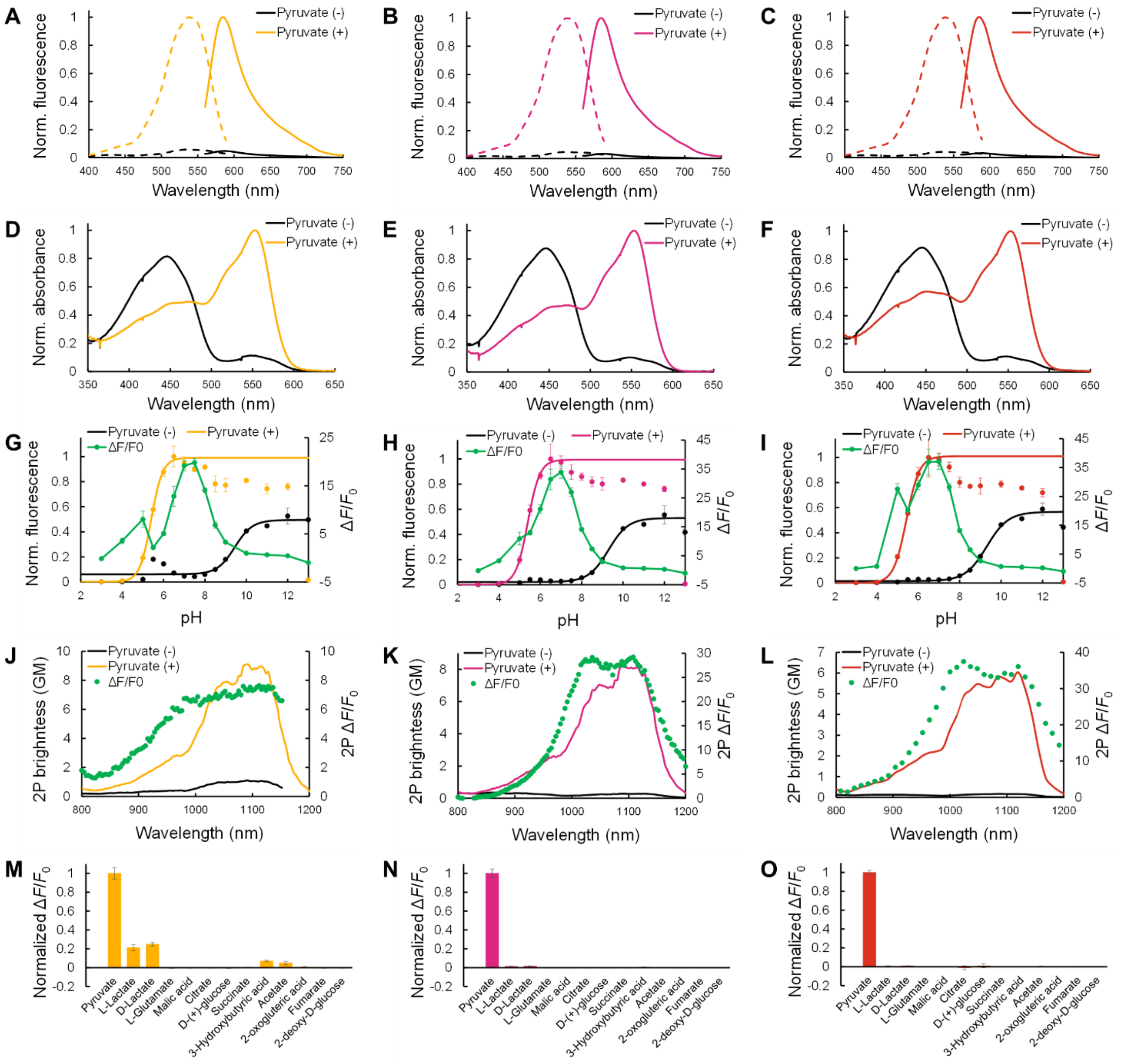

**Figure S8.** ***In vitro* characterization of ApplePy affinity variants**

**A-C,** Excitation (emission at 640 nm) and emission (excitation at 510 nm) spectra of ApplePy1Highest (**A**), ApplePy1Low (**B**) and ApplePy1Lowest (**C**) in the presence (100 mM) and absence of pyruvate. **D-F,** Absorbance spectra of ApplePy1Highest (**D**), ApplePy1Low (**E**) and ApplePy1Lowest (**F**) in the presence (10 mM) and absence of pyruvate. **G-I,** pH titration curve of ApplePy1Highest (**G**), ApplePy1Low (**H**) and ApplePy1Lowest (**I**) in the presence (100 mM) and the absence of pyruvate (n = 3 replicates; mean ± s.d.). The curve was fitted to calculate the physiological p$K_a$ value. **J-L,** Two-photon excitation spectra of ApplePy1Highest (**J**), ApplePy1Low (**K**) and ApplePy1Lowest (**L**) in the presence (10 mM) and absence of pyruvate. 2P, Two-photon. GM, Goeppert-Mayer units. **M-O,** Fluorescence responses of ApplePy1Highest (**M**), ApplePy1Low (**N**) and ApplePy1Lowest (**O**) to various metabolites at 10 mM (n = 3 replicates; mean ± s.d.).

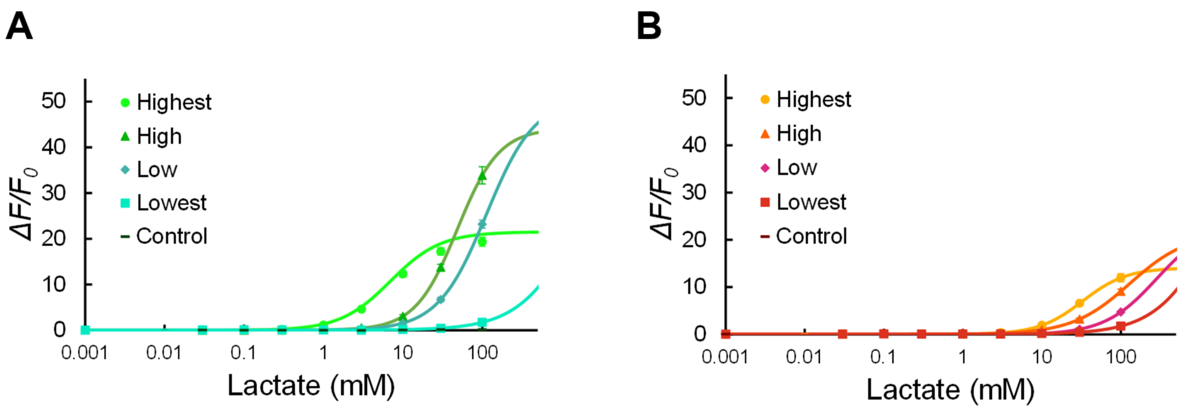

**Figure S9. Selectivity toward L-lactate.**

**A,B,** L-lactate titration curve of purified GreenPy1 (**A**) and ApplePy1 (**B**) variants (n = 3 replicates; mean ± s.d.).

#### Supplementary References

1. Hario, S. *et al.* High-Performance Genetically Encoded Green Fluorescent Biosensors for Intracellular l-Lactate. *ACS Cent Sci* **10**, 402–416 (2024).
2. Kalivoda, K. A., Steenbergen, S. M. & Vimr, E. R. Control of the Escherichia coli sialoregulon by transcriptional repressor NanR. *J. Bacteriol.* **195**, 4689–4701 (2013).
3. Condemine, G., Berrier, C., Plumbridge, J. & Ghazi, A. Function and expression of an N-acetylneuraminic acid-inducible outer membrane channel in Escherichia coli. *J. Bacteriol.* **187**, 1959–1965 (2005).
4. Suvorova, I. A., Korostelev, Y. D. & Gelfand, M. S. GntR family of bacterial transcription factors and their DNA binding motifs: Structure, positioning and co-evolution. *PLoS One* **10**, e0132618 (2015).
